# Aberrant splicing in Huntington’s disease via disrupted TDP-43 activity accompanied by altered m6A RNA modification

**DOI:** 10.1101/2023.10.31.565004

**Authors:** Thai B. Nguyen, Ricardo Miramontes, Carlos Chillon-Marinas, Roy Maimon, Sonia Vazquez-Sanchez, Alice L. Lau, Nicolette R. McClure, Whitney E. England, Monika Singha, Jennifer T. Stocksdale, Ki-Hong Jang, Sunhee Jung, Jharrayne I. McKnight, Leanne N. Ho, Richard L.M. Faull, Joan S. Steffan, Jack C. Reidling, Cholsoon Jang, Gina Lee, Don W. Cleveland, Clotilde Lagier-Tourenne, Robert C. Spitale, Leslie M. Thompson

## Abstract

Huntington’s disease (HD) is a neurodegenerative disorder caused by a CAG repeat expansion in the first exon of the *HTT* gene encoding huntingtin. Prior reports have established a correlation between CAG expanded *HTT* and altered gene expression. However, the mechanisms leading to disruption of RNA processing in HD remain unclear. Here, our analysis of the reported HTT protein interactome identifies interactions with known RNA-binding proteins (RBPs). Total, long-read sequencing and targeted RASL-seq of RNAs from cortex and striatum of the HD mouse model R6/2 reveals increased exon skipping which is confirmed in Q150 and Q175 knock-in mice and in HD human brain. We identify the RBP TDP-43 and the N6-methyladenosine (m6A) writer protein methyltransferase 3 (METTL3) to be upstream regulators of exon skipping in HD. Along with this novel mechanistic insight, we observe decreased nuclear localization of TDP-43 and cytoplasmic accumulation of phosphorylated TDP-43 in HD mice and human brain. In addition, TDP-43 co-localizes with HTT in human HD brain forming novel nuclear aggregate-like bodies distinct from mutant HTT inclusions or previously observed TDP-43 pathologies. Binding of TDP-43 onto RNAs encoding HD-associated differentially expressed and aberrantly spliced genes is decreased. Finally, m6A RNA modification is reduced on RNAs abnormally expressed in striatum from HD R6/2 mouse brain, including at clustered sites adjacent to TDP-43 binding sites. Our evidence supports TDP-43 loss of function coupled with altered m6A modification as a novel mechanism underlying alternative splicing/unannotated exon usage in HD and highlights the critical nature of TDP-43 function across multiple neurodegenerative diseases.

## Main Text

Huntington’s disease (HD) is an autosomal dominant neurodegenerative disorder that manifests with motor, cognitive, and psychiatric symptoms^1–3^. HD is caused by a CAG repeat expansion mutation in exon 1 of the Huntingtin (*HTT*) gene, which encodes an expanded polyglutamine repeat within the Huntingtin (HTT) protein (mHTT)^1^. The most overt pathological phenotypes include loss of striatal neurons, primarily the GABA-ergic medium spiny neurons, and cortical atrophy^4^. Somatic repeat instability occurs in striatum^5–9^ and the mutation is linked to both gain of pathogenic functions and disruption of normal HTT function^10^. The CAG expansion mutation also promotes the formation of an aberrant and toxic splice product through incomplete splicing of exon 1, mHTT ^11^, and accumulation of repeat associated non-ATG (RAN) translation proteins^12^. To date, no disease-modifying treatment is available.

HTT protein interactors and how those interactions change in the context of mHTT expression have been documented in yeast, mammalian cells, mouse brain, and human brain^13^. One class of HTT-protein interactors is represented by RNA-binding proteins (RBPs), which are responsible for all aspects of RNA processing^14,15^. Of the published work on HTT’s and RBPs, the majority [e.g. FUS^16^, G3BP1^17,18^, CAPRIN1^17^, VCP^19^ and TDP-43^18,20,21^ have been linked to regulation of the oxidative stress response and aggregation. While abnormal interactions between HTT and RBPs were reported, suggesting roles in RNA processing, the mechanisms by which mHTT leads to alterations of RNA expression and splicing—a hallmark of HD and other neuropathological disorders—remain undetermined.

TDP-43 is a DNA/RNA binding protein critical for splicing regulation that is mutated and/or mis-localized in amyotrophic lateral sclerosis (ALS) and frontotemporal lobar degeneration (FLTD)^22–24^. Indeed, a striking nuclear to cytoplasmic translocation of TDP-43 is the pathological hallmark in 97% of sporadic ALS (sALS) cases and up to 50% of sporadic and familial FTLD patients^22,25–27^. TDP-43 mis-localization is also found in most Alzheimer’s disease (AD) cases^28^, a subset of patients with Parkinson’s Disease^29,30^, and in HD patient brain^20,31^. In HD, TDP-43 can co-localize with HTT EM48 positive intracellular inclusions^20^, and was found to promote somatic CAG repeat expansion in HD mouse model^18^. Loss of nuclear TDP-43 results in abnormal splicing events^23,32,33^ including the aberrant inclusion of cryptic exons^34–38^. It is not known whether HD-associated expression and splicing alterations are related to TDP-43 disruption in HD.

The most abundant chemical modification of mRNA—m6A-methylation (m6A)—occurs on the adenosine base at the N6 position, altering localization, splicing, stability, and translational control^39–49^. Adenosine is methylated to yield m6A by writer proteins (METTL3, METTL14, WTAP, RBM15), and reversed by eraser proteins (FTO, ALKBH5). m6A attracts and repels RBPs to modulate the binding and regulation of the target RNA^50^, and regulates neural development^46,51^. It was recently shown that post-mortem tissues from ALS patients display widespread RNA hypermethylation and that TDP-43 preferentially binds m6A-modified RNAs^52^. Additionally, altered m6A patterns have been reported in the hippocampus of the Q111 knock-in mouse model of HD with effects on behavior^53^.

Here, RNA-seq analysis of brain tissue from the HD R6/2 model^54^ that expresses transgenic human mHTT exon1 was used to identify alterations in RNA processing including RNA splicing. Splicing-specific changes were also identified in HD Q150 and Q175 knock-in mouse models. Novel processing alterations were confirmed by long-read RNA sequencing. We identified primary sequence motifs—RNA binding sites associated with mHTT-dependent changes in alternative splicing using primary sequence models—that implicated two HTT-interacting RBPs: TDP-43 and Methyltransferase 3 (METTL3). Using molecular and neuropathological measures and TDP-43 CLIP-seq and m6A CLIP-seq, we determined that mHTT disrupts TDP-43 and METTL3 function in post-transcriptional processing of their RNA targets in HD. We further show a novel nuclear aggregate-like structure in HD patient brain that contains TDP-43 and HTT. This study provides the first evidence for functional disruption of TDP-43 in HD and a novel association with abnormal m6A RNA modification in HD. Further, this work highlights that TDP-43 dysregulation may be a fundamental driver of pathogenesis in a broader group of diseases than previously appreciated and suggests that biomarker development and therapeutic approaches focused on TDP-43 may represent fundamental strategies across neurodegenerative diseases.

## Results

### RNA processing dysregulation in striatum and cortex of HD mice

HTT and mHTT protein-protein interactors have been identified from human, mouse, cell, and yeast and were curated into a list of 4,182 predicted protein interactors (HdinHD.org)^13^. We found that the top enriched Gene Ontology (GO) function term was RNA binding, with 788 interactors (18.8%) being known or predicted RBPs (42.9% (788/1836)^55^ (Fig. 1A, RBP interactors of HTT table S1). RNA-sequencing was performed to analyze splicing alterations in ten 3-month-old (3mos) mice per group (5 males, 5 females; HD vs NT) symptomatic HD R6/2 transgenic and non-transgenic (NT) control mice^54^ considering this model’s transcriptional signature that recapitulates many human HD signatures^56^. Principal Component Analysis (PCA) showed separation of genotypes across PC1 and sex across PC2 (Fig. S1A). Differential Gene Expression (DEG) analysis accounting for sex differences resulted in 1,309 downregulated genes and 648 upregulated genes in the striatum (adjusted P-value of < 0.05 and log2foldchange > 1), and 1,253 downregulated genes and 462 upregulated genes in the cortex (significant DEGs with adjusted P-value of < 0.05 in table S2). A ∼65.8% overlap between the striatal and cortical differentially expressed genes was observed with 95.4% changing in the same direction (Fig. S1B). A HD striatal transcriptomic signature of 266 disease-associated expression changes was previously identified from multiple mouse models of HD (HD266ssg)^57^. Our striatal RNA-seq results showed dysregulation of all 266; 265 in the anticipated direction. The cortical transcriptome signature generally mirrored the striatal 266 genes (Fig. S1C, full 266 gene list in table S3). Strengthening our hypothesis that HTT’s interactions with RBPs may drive HD transcriptional signatures in the cortex and striatum, we observed up-regulated DEGs in both regions to be enriched for gene ontology terms relating to RNA metabolism, RNA processing, and RNA splicing (Fig. 1B & 1C).

**Figure 1:**
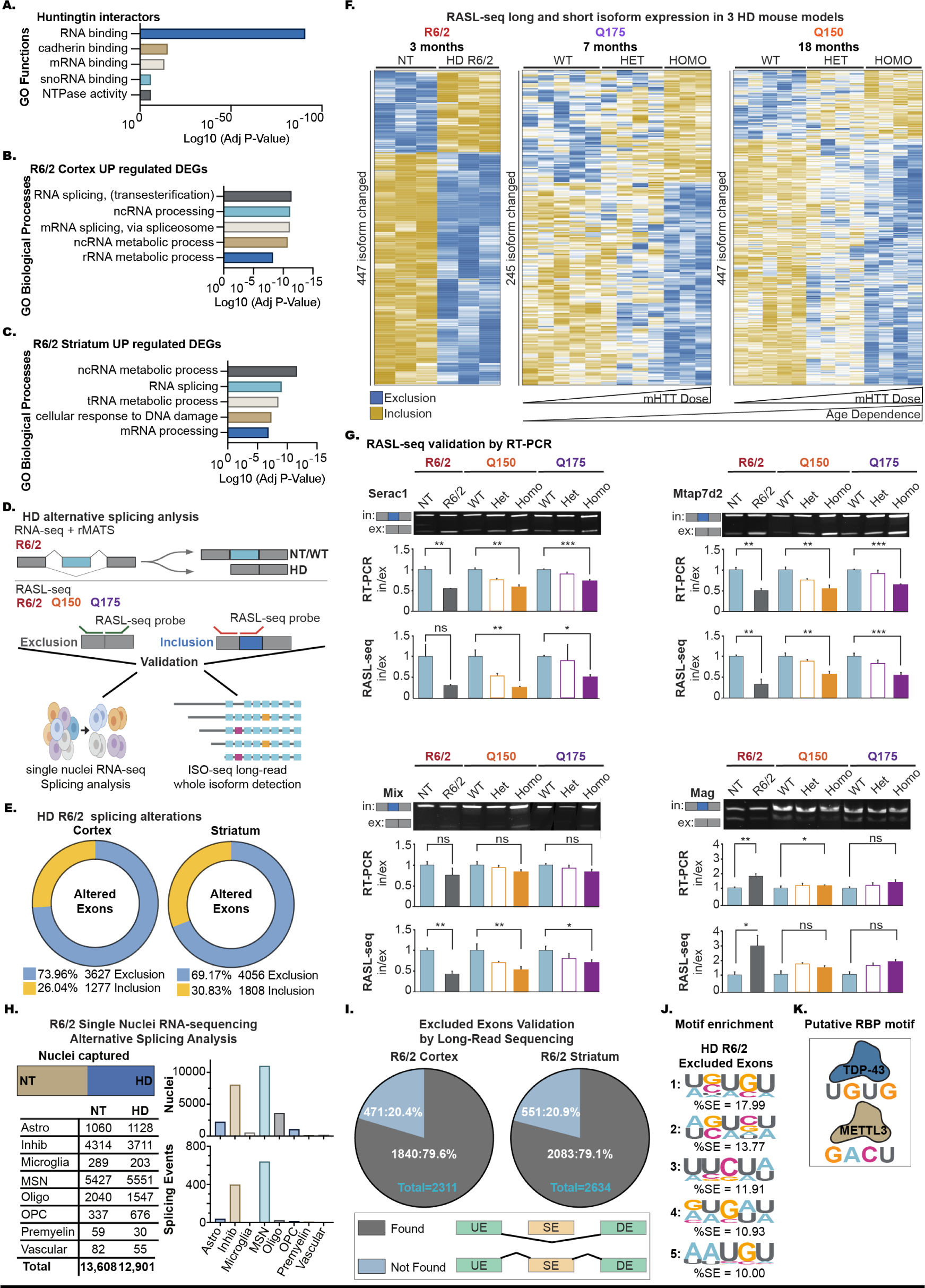
Dysregulated RNA processing in HD R6/2 mouse striatum and cortex. (A) Gene ontology analysis for molecular functions of 4182 predicted protein interactors of Huntingtin ^13^. X-axis represents the log10(adjusted p-value). Gene ontology analysis for biological processes of upregulated genes identified by DEseq2 in 3 months old R6/2 mice from cortex (B) and (C) striatum. X-axes represent the log10(adjusted p-value). (D) Schematics of detection of alternatively spliced exons in HD mouse models with RASL-seq and rMATS. (E) Alternative splicing events annotated by rMATS: skipped exons detected in cortex (left) and striatum (right) from 3 months old R6/2. Splicing events filtered for FDR < 0.05. (F) Heatmap showing the detection of the short isoform (skipped exon) and long isoform (included exon) from RASL-seq in cortex from the R6/2, Q175, Q150 HD mouse models. NT – Non transgenic control for the R6/2, WT – Wildtype, HET – Q7/Q175 – Q7/Q150, HOMO – Q175/Q175 – Q150/Q150. (G) PCR primers were generated to validate the included and excluded isoforms detected by RASL-seq. (H) Single nuclei sequencing validation; (left) cell type classification of total nuclei sequenced from the striatum of 12wks R6/2. From top to bottom; Astro = Astrocytes, Inhib = inhibiatory neurons, Microglia, MSN = Medium Spiny Neurons, Oligo = Oligodendrocytes, OPC = Oligodendrocyte progenitor cells, Premyelin cells, & Vascular cells. (right) graph shows the amount of altered splicing events detected per cell type (1,109 total). (I) Pie chart showing the detection of significant skipped exons in PacBio ISO-seq long-read sequencing. (J) Top 5 enriched 5-mer binding motifs from de novo motif enrichment by HOMER of the HD-associated skipped exons. (K) Canonical binding motif for the TDP-43 (UG rich), and METTL3 (DRACH).

### Splicing defects in HD mice are characterized by increased exon exclusion

Next, we set out to understand alternative splicing (AS) changes in the HD mouse models (Fig. 1D). Applying the AS tool (RNA-seq Multivariate Analysis of Transcript Splicing (rMATS)^58^) on R6/2 and age-matched NT RNA-seq data from 3mos animals, 9,033 RNAs (3,620 genes) with significant splicing events (FDR < 0.05) were identified in the cortex and 10,074 RNAs (3,854 genes) in the striatum (Fig. S1D left). These are classified by rMATS into 5 canonical splicing events: skipped cassette exons (SE), alternative 5’ splice sites (A5SS), alternative 3’ splice sites (A3SS), mutually exclusive exons (MXE), and retained introns (RI) (Fig. S1D right). Segmentation of the significant AS changes by condition (HD vs. NT Control) revealed ∼70% of SE events result in higher exon exclusion in HD in both affected brain regions, cortex and striatum (Fig. 1E, table S4). GO analysis found an enrichment for abnormal exon exclusion in cortical and striatal genes responsible for synaptic transmissions in neurons (Fig. S1E & S1F). Consistent with AS events contributing to gene expression alteration, ∼60% of the genes with AS are up- or down-regulated in cortex and striatum of R6/2 mice (Fig. S1G). We then employed a splicing-specific sequencing strategy (<u>R</u>NA-mediated oligonucleotide <u>A</u>nnealing, <u>S</u>election, and <u>L</u>igation with next-generation sequencing; RASL-seq)^59,60^ with 9,496 primer pairs targeting known splice junctions in the cortex of HD R6/2 and two additional knock-in HD mouse models expressing 150 repeats (Q150)^61^ or 175 repeats (Q175)^61^ at 3, 7, and 18 months, respectively. Consistent with our findings from RNA-seq and rMATS analysis, RASL-seq identified an increase in exon exclusion events in 3mos R6/2 compared to NT littermates (Fig. 1F left). Furthermore, we observe an mHTT dose-dependent and age-dependent increase in exon exclusion in the Q150 and Q175 mice (Fig. 1F middle and right). Validation of splicing alterations identified by RASL-seq was performed using semi-quantitative RT-PCR for genes that overlapped (Serac1, Mtap7d2, Mlx, Mag) between the three mouse models (Fig. 1G & S1H).

### Long-read single transcript RNA sequencing and single nuclei RNA sequencing validates splicing events in HD R6/2

We validated the increase in HD exon exclusion events using single nuclei RNA-seq and long read sequencing (Iso-seq)^62^. The RNA-seq data does not account for the cellular heterogeneity of the striatum; therefore, we determined whether there is a cell-type specificity for the observed splicing signature. We previously carried out single nuclei RNA-seq on the striatum and cortex from a different cohort of male HD R6/2 and NT animals at the same symptomatic 3mos stage^63^. Splicing analysis of this dataset using DESJ-detection ^64^ showed that most of the splicing alterations were arising in the inhibitory medium spiny neurons, the cell type most affected in HD (Fig. 1H). We also employed long-read sequencing using the Pacific Biosciences iso-seq platform as a high-throughput method to validate our detected splicing changes by sequencing full-length mRNA-enriched transcripts as described^62^. From a subset of the samples described above for short-read RNA-seq (n=4 per condition, males only), we captured 56,000 isoforms from the cortex and 53,000 isoforms from the striatum. Because we detected the most significant AS differences for exon exclusion, we focused our analysis on this splicing event. Long-read sequencing identified 1840/2311 (79.6%) genes with significant exon exclusions detected by short-read sequencing in the cortex of R6/2 mice (Fig. 1I left) and 2083/2634 (79.1%) significant exon exclusions in the striatum (Fig. 1I right).

### *De Novo* motif analysis of HD skipped exons reveals an enrichment for the binding motif of TDP-43 and the RNA modification N6-methyladenosine (m6A)

Since an enriched number of HTT interactors are RBPs, we investigated whether the presence of mHTT may alter RNA splicing by disrupting RBP interactions with their RNA targets. We used HOMER^65^ to perform a *de novo* motif analysis for the excluded exons affected in HD. The analysis revealed significant but modest enrichment for the UGUGU and UGACU RNA sequence motifs (Fig. 1J). UG/GU-rich motifs are RBP preference sites for TAR DNA-Binding Protein (TARDBP/TDP-43)^23,32,66^. In addition, HOMER analysis also revealed that the second potential regulatory binding motif is the canonical N6-methyladenosine (m6A) DRACH motif (D=A/G/U, R=A/G, H=A/C/U; including UGACU) recognized by writer proteins such as METTL3^67,68^ (Fig. 1K). We investigated whether alterations in these two proteins could contribute to the RNA processing alterations in the HD mice. Loss of TDP-43 is associated with downregulation of long intron-containing genes^23,24^. Consistent with a disruption of TDP-43 in HD mice, transcripts downregulated in the cortex and striatum of R6/2 mice have, on average, longer introns than upregulated genes (Fig. S2A). TDP-43 autoregulates its own transcript by binding its 3’UTR and loss of nuclear TDP-43 protein results in a modest increase of TDP-43 mRNAs^69^. Analysis of TDP-43 RNA expression level by RNA-seq in R6/2 mice and by RT-qPCR in R6/1 mice (Fig. S2B & S2C) reveals a modest increase in TDP-43 RNA levels, supporting a loss of nuclear TDP-43. We next compared the HD cortical and striatal excluded exons to splicing alterations identified in primary neurons from wild-type mice with antisense oligonucleotides (ASO)-mediated depletion of TDP-43^70^ and observe a 43.5% gene overlap in dysregulated, with ∼50% in the same direction (Fig. S2D). Our data suggests that TDP-43 contributes to transcriptional dysregulation in HD that may involve an exciting intersection between TDP-43 and m6A in HD that has not been previously described. Therefore, we investigated how TDP-43 and the m6A RNA modification might contribute to altered splicing and HD pathology.

### Decreased nuclear expression of TDP-43 in brains from HD mouse models and HD patients

TDP-43 is one of the RBPs listed as an interactor of HTT protein^13,20,21^ including by proteomic analysis^68^. We confirmed this interaction by immunoprecipitation of TDP-43 from mouse brain with mouse full-length HTT (FL-HTT). Notably, the interaction between TDP-43 and mHTT is reduced in cortex of R6/2 mice compared to NT mice (Fig. S2E). We assessed TDP-43’s protein expression by immunofluorescence (IF) in cortex from 3mos late symptomatic R6/2 mice and observed decreased nuclear TDP-43 protein expression. Similar TDP-43 nuclear reduction was observed in cortex from 11mos R6/1 mice and 24mos homozygous Q175 mice (Fig. 2A & Fig. S2G). This reduced nuclear staining of TDP-43 did not result from reduced TDP-43 protein levels, as quantitative western blot analysis showed equivalent TDP-43 levels in HD R6/2 cortex (Fig. S2F). This decrease of nuclear TDP-43 was also found in human HD postmortem superior frontal gyrus (SFG) compared to tissue from unaffected individuals by immunofluorescence staining (Control: n=2, HD: n=5) (Fig. 2B & Fig. S2H). Automated quantification of nuclear TDP-43 levels using CellProfiler revealed a significant decrease in nuclear TDP-43 fluorescent intensity in all 5 HD compared to control tissues (Fig. 2C), strongly supporting loss of TDP-43 function as a contributor to HD pathogenesis.

**Figure 2:**
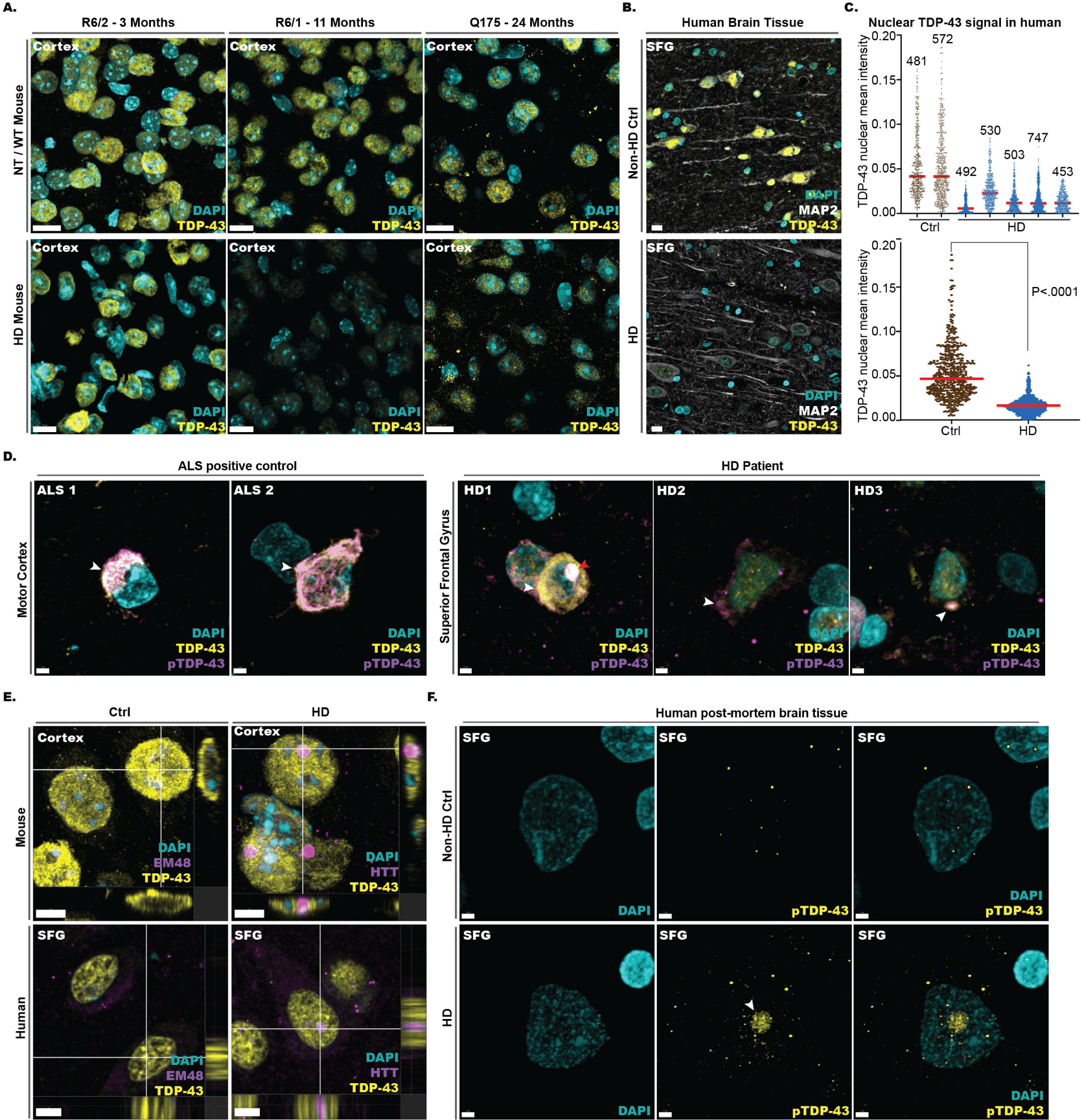
HD systems revealed decreased TDP-43 nuclear expression, increased phosphorylated-TDP-43 accumulation in the cytoplasm, and Aggregation-Like phosphorylated TDP-43 nuclear Bodies. (A) Immunofluorescence (IF) staining of the cortex of 3-month-old HD R6/2 mice (left), 11-month-old HD R6/1 mice (middle), 24-month-old Homozygous Q175 mice (right) and wild-type littermates (NT or WT). Cyan is the nuclei marker DAPI, yellow shows total TDP-43 staining. (B) Representative IF staining images of superior frontal gyrus from HD patients compared to non-HD control individuals showing decreased TDP-43 (yellow) signal intensity. (C) Quantification of decreased nuclear TDP-43 signal intensity, 5 representative images were taken at 40X from 5 HD and 2 control individuals. A Cell profiler pipeline was created to identify larger nuclei (enriched for neurons) by dapi staining. The average of TPD-43 nuclear signal was obtained by measuring the intensity signal within a mask defined by DAPI. Each cell’s mean nuclear TDP-43 intensity is plotted. One-way ANOVA was performed with multiple comparisons and resulted with significant changes between all HD vs Ctrl comparisons. Unpaired t-test between control vs HD (bottom) p-value < 0.0001. Numbers on top of each group indicate the number of cells plotted. Red bar represents the medium. (D) Representative IF staining images of the motor cortex from an ALS patient (positive control) compared to HD patients, using antibodies against total TDP-43 (yellow), phosphorylated TDP-43 (purple), and nuclear stain DAPI. White arrow indicates pTDP-43 cytoplasmic aggregation. Red arrow (HD1), cytoplasmic aggregate verified by orthogonal view. (E) Orthogonal view of TDP-43’s co-localization with mHTT nuclear inclusion stained with EM48 (left) in 3-month HD R6/2 and NT, and human HD postmortem brain tissue and non-HD control (right). (F) Representative images of immunofluorescence (IF) staining of pTDP-43 (yellow) nuclear AL-bodies identified in superior frontal gyrus (SFG) of HD patients and not in unaffected control individuals.

### Detection of cytoplasmic phosphorylated TDP-43 and co-localization of TDP-43 with HTT nuclear inclusions in HD patient post-mortem brains

To examine nuclear-cytoplasmic translocation and phosphorylation of TDP-43 (pTDP-43), we co-stained brain tissues from HD and unaffected individuals with a TDP-43 antibody and a pTDP-43 antibody specific to phosphorylated serine 409 and serine 410 (S409/S410). We detected, by IF, the presence of rare cytoplasmic TDP-43 aggregates, recognized by both antibodies, in Map2-positive neurons of HD patients but not in unaffected individuals (Fig. 2D). We observed an aggregate-like morphology in the perinuclear space, similar to but less dense and fibrous than, pTDP-43 IF in ALS/FTD^22,25–27^ or in ALS motor cortex positive controls evaluated (Fig. 2D, low magnification Fig. S3A, separated channels Fig. S3B).

We next evaluated whether TDP-43 colocalized with mHTT nuclear inclusions by co-staining for TDP-43 and mHTT using two independent antibodies (EM48 and MW8) that detect mHTT inclusions^71^ in cortex and striatum from 3mos R6/2 mice. We observed diffuse nuclear TDP-43 distribution and modest co-localization to mHTT nuclear inclusions in a subset of neurons (Fig. 2E top, Imaris co-localization histogram Fig. S3C). In superior frontal gyrus (SFG) from HD patients (HD=6, CTRL=6), HTT nuclear inclusions were detected in ∼10%-20% of the neurons using an N-terminus HTT antibody (MAB5492 & EPR5526) and modest TDP-43 co-localization with these inclusions (Fig. 2E bottom, Fig. S3D). Despite the nuclear reduction of TDP-43 in HD mice and human, a subset of neurons showed co-localization with nuclear HTT inclusions. Our results suggest that the presence of mHTTex1 causes mis-localization of TDP-43, including within nuclear HTT inclusions, which could alter its normal nuclear function.

### Nuclear accumulation of phosphorylated TDP-43 with HTT in aggregation-like bodies (AL-bodies)

In addition to cytoplasmic accumulation of pTDP-43, and co-localization of TDP-43 with nuclear HTT inclusions, we observed an, as of yet, not previously described spherical accumulation of fibrous-like pTDP-43 in the nucleus. These structures measure an average of ∼3 microns in diameter and are not detectable in the control human tissues (Fig. 2F). We refer to these structures as pTDP-43 nuclear Aggregation-Like bodies (AL-bodies). pTDP-43 AL-bodies colocalize with n-terminal TDP-43 (Fig. S4A) and were also detected with a second S409/S410 pTDP-43 antibody (Fig. S4B (low magnification) & S4C (high magnification)). We observed these AL-bodies in the majority of Map2-positive neurons (Fig. S4D) and detected co-localization of AL-bodies and HTT (5526) by IF in HD patient tissue (Fig. S4E). Notably, the co-localization of pTDP-43 and HTT protein in AL-bodies is diminished in cells with canonical HTT nuclear inclusions (Fig. S4F. This striking accumulation of nuclear pTDP-43 and HTT into distinct spherical aggregation-like bodies in MAP2-positive neurons from HD patient brains represents a novel type of TDP-43 pathology that may be unique to HD.

### TDP-43 binding is attenuated in dysregulated HD genes

To further investigate TDP-43’s impact on transcriptional regulation in HD, we focused on RNA targets of TDP-43 by reanalyzing published TDP-43 CLIP-seq data (Fig. S5A) from WT mouse striatum^23^. We remapped the TDP-43 CLIP-seq data to the mm10 reference genome and recapitulated the UGUGU binding motif (Fig. S5B). As expected, TDP-43 binds to 79/266 (29.7%) HD266ss genes (Fig. S5C). We reasoned that if TDP-43 is a central regulator of HD gene expression, then KD by an ASO should capture at least partial gene expression changes within the HD266ssg. Therefore, from the same dataset, we reprocessed RNA-seq data from mouse striatum with TDP-43 ASO or control ASO treated, age-matched R6/2 and NT data from our published report^72^. 2/4 TDP-43 ASO treated striatum clustered with the HD R6/2 animals on a gene expression heatmap of the 79/266 HD266ssg (Fig. 3A), with trending gene expression changes for established HD genes Pde10a, Adora2a, Homer1, and Drd2 (Fig. 3B). We next tested whether TDP-43 knockdown by siRNA in isogenic induced pluripotent stem cell (iPSCs) lines expressing nonpathogenic length HTT-18Q or HD causing mHTT-50Q and differentiated to medium spiny neurons (MSNs)^73^, would show similar patterns. After confirmation of TDP-43 protein KD by western blot analysis (fig 3D), samples were subjected to RNA-seq. Our analysis revealed 4,870 DEGs (3,121 down regulated, 1,749 up regulated) when comparing HTT-18Q versus mHTT-50Q (table S5) – “*genotype”* effect. For the “*TDP-43 KD”* effect, we observed 951 DEGs (556 down regulated, 395 up regulated), comparing HTT-18Q TDP-43 KD vs scramble siRNA control (table S6). We applied the hypergeometric test and observed with significance that 68.7% (382/556, *P*=1.5x10^-189^) of the TDP-43 KD down regulated DEGs overlap with HD down regulated DEGs, and 46.3% (183/395, *P*=7.5x10^-93^) overlap with the HD up regulated DEGs (Fig. 3E, table S7). Notably, we observe STMN2, UNC13B, and CAMK2B as downregulated DEGs in both HD and TDP-43 KD comparisons (Fig. 3F). Together our analysis suggests that TDP-43 KD drives similar gene expression pattern changes as mHTT in HD MSNs.

**Figure 3:**
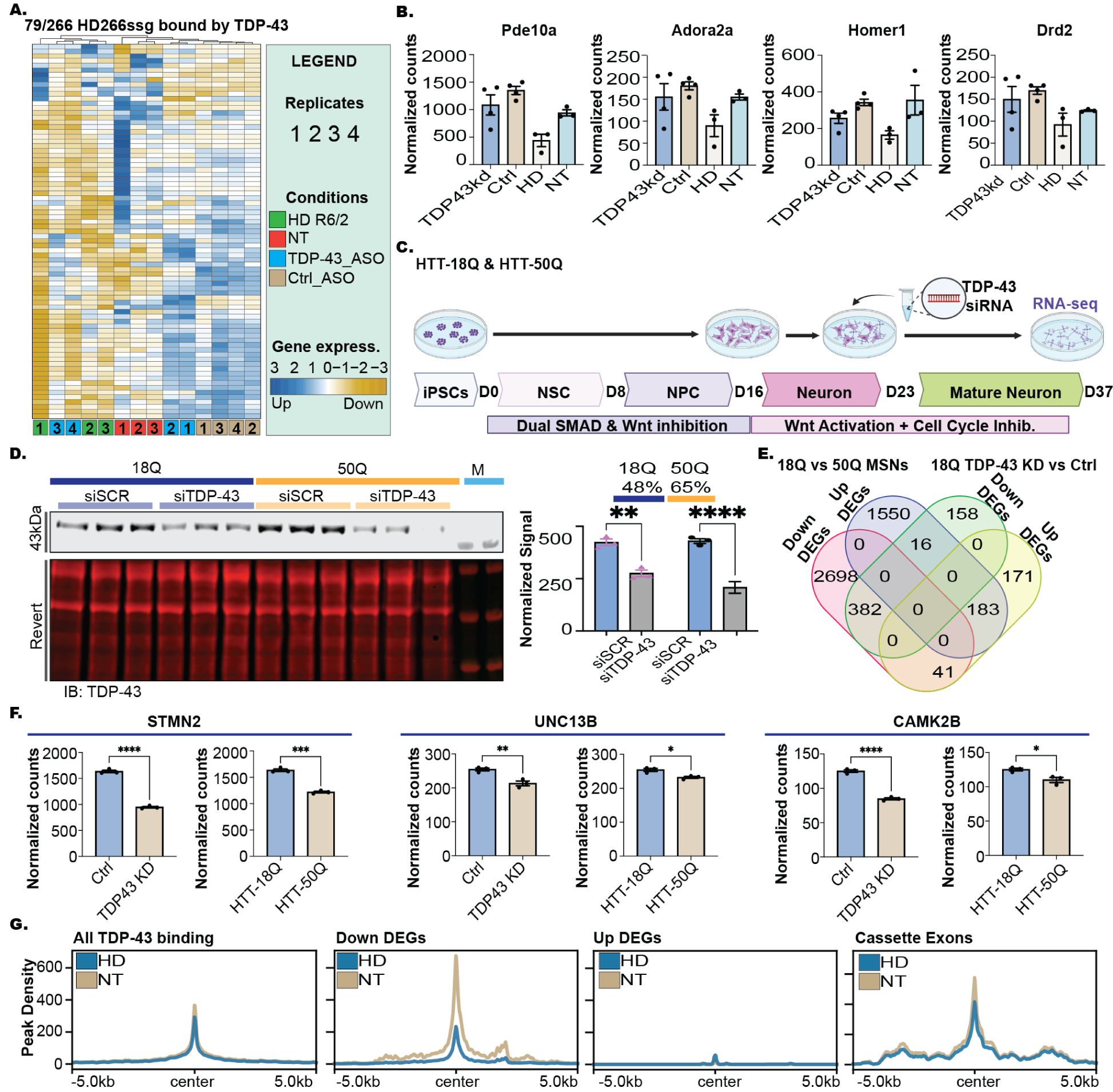
TDP-43 changes in Huntington’s disease. (A) Heatmap showing clustering of 8wks HD R6/2, NT, TDP-43 ASO treated, and Control ASO treated on gene expression of the 79/266 HD striatal genes that are RNA targets of TDP-43. (B) Normalized counts from RNA-seq data for signature HD striatal genes; Pde10a, Adora2a, Homer1, and Drd2. (C) Schematic of iPSC differentiation into medium spiny neurons with TDP-43 KD by siRNA. (D) Western blot for TDP-43 protein levels after treatment of MSNs with TDP-43 siRNA. Right bar graph plots TDP-43 intensity normalized to Revert total protein stain. Stastical significance was determined by un-paired T-test. (E) Venn Diagram showing the overlap of DEGs between HTT-18Q MSNs scramble control vs. TDP-43 siRNA and HTT-18Q MSNs vs mHTT-50Q. (F) Example of key gene expression changes anticipated from TDP-43 KD. (G) Reproducible R6/2 HD and NT IDR TDP-43 peaks were centered and plotted on all TDP-43 binding sites, compared to Down DEGs, Up DEGs, and Cassette Exons.

To identify TDP-43 RNA targets that may be altered by the presence of mHTTex1, we also carried out TDP-43 enhanced Cross-Linking Immuno-Precipitation followed by sequencing (eCLIP-seq^74^) on HD R6/2 and NT mouse cortex and striatum (n= 8, age= 3 mos, 4 males, 4 females). Surprisingly, our binding site distribution primarily spanned coding regions and 3’UTR, unlike previous findings^23,32^ that showed primarily intronic binding (Fig. S5D). This difference could be due to regional specificity (striatum/cortex versus whole brain), or a technical aspect of our experiment which enriched for processed mRNAs. Analysis of 5mer U-rich motifs in our TDP-43 CLIP peaks reveals an enrichment for UGUGU as previously reported (Fig. S5E). The TDP-43 eCLIP-seq yielded ∼60% target gene overlap with our reanalysis of Polymenidou et. al, 2011^23^. There is an ∼80% overlap between HD and NT TDP-43 binding to RNA transcripts in both brain regions, suggesting that most targets remain unchanged due to mHTTex1 expression (Fig. S5G). Mapping TDP-43 significant peaks to genes with excluded exons identified from HD R6/2 cortex and striatum (Fig. S2A) revealed TDP-43 binding on ∼57-64% of genes containing exon exclusions (Fig. S5H). Next, we assessed the possibility that TDP-43 drives HD transcriptional dysregulation through direct binding to genes constituting the striatal HD signature. TDP-43 binds ∼30% of HD266ssg with the majority of resulting in downregulation of transcripts. Plotting TDP-43 binding density across target genes, HD down regulated, HD up regulated, and HD cassette exons, TDP-43 binding is decreased on down regulated genes in HD. There is also slightly decreased binding on cassette exons which includes excluded and included exons (Fig. 3G); however, upregulated genes do not show the same pattern. This suggests that loss of TDP-43 binding to target genes can decrease gene expression of the target genes and promote cassette exon splicing. Together our observations reveal that decreased TDP-43 binding is specific to dysregulated HD striatal genes and drives HD transcriptomic signature.

### m6A-sequencing reveals an altered epitranscriptome in the HD R6/2 cortex and striatum

The second potential regulatory binding motif from the RNA-seq study is the canonical m6A motif (Fig. 1G). To further understand how the HD epitranscriptome may contribute to HD and observed alternative splicing changes, we performed immunofluorescence staining for the proteins of the m6A machinery, both m6A writers (METTL3, METTL14, RBM15, WTAP) and erasers (ALKBH5, FTO), in the cortex and striatum from HD R6/2 at 3mos (late symptomatic) of age (n=8, males and females). Since the m6A machinery proteins are localized to the nucleus, we used Cell Profiler to segment out the nuclear signal and quantitated the average intensity per cell. Applying the same methodology across all the stained sections, a significant decrease in nuclear expression was identified for METTL3 in both the cortex and striatum (Fig. 4A & 4B) and for RBM15 in the cortex, while the other m6A machinery proteins were unchanged (Fig. S5I). With a reduction in METTL3 protein expression, we expect m6A levels to be decreased in the HD R6/2 mice. However, liquid chromatography-mass spectrometry (LC-MS) analysis of polyA-selected RNA from mouse cortex (n=10 males & females) samples did not show a significant global change of m6A levels (Fig. 4C). Because the global LC-MS analysis cannot capture m6A modification changes in a subset of genes, we carried out single nucleotide resolution transcriptome-wide m6A sequencing using eCLIP-seq with an m6A-specific antibody^74^. m6A eCLIP-seq identified 28,8976 shared, 5,273 HD, and 3,035 NT m6A genomic locations in the cortex and 30,459 shared, 3,355 HD, and 3,905 NT in the striata, respectively (Fig. 4D). Motif enrichment revealed that >70% of the detected m6A peaks fall within the expected DRACH m6A motif as previously reported (Fig. S5J) and recapitulate the concentrated distribution of m6A sites at the 3’UTR in a metagene plot (Fig. S5K). Plotting the m6A sites found on HD DEGs across a metagene plot revealed an altered epitranscriptomic landscape in HD R6/2 mice (Fig. 4E). We then determined whether m6A deposition can influence the HD transcriptome by evaluating enrichment for m6A deposition on HD DEGs. After normalizing for length, DEGs have significantly more m6A sites than non-DEGs (Fig. 4F), and within the DEGs, the genes that go on to be downregulated are enriched for number of m6A sites (Fig. 4G). Overall, these data suggest that m6A deposition is important for the steady-state expression and processing of HD DEGs.

**Figure 4:**
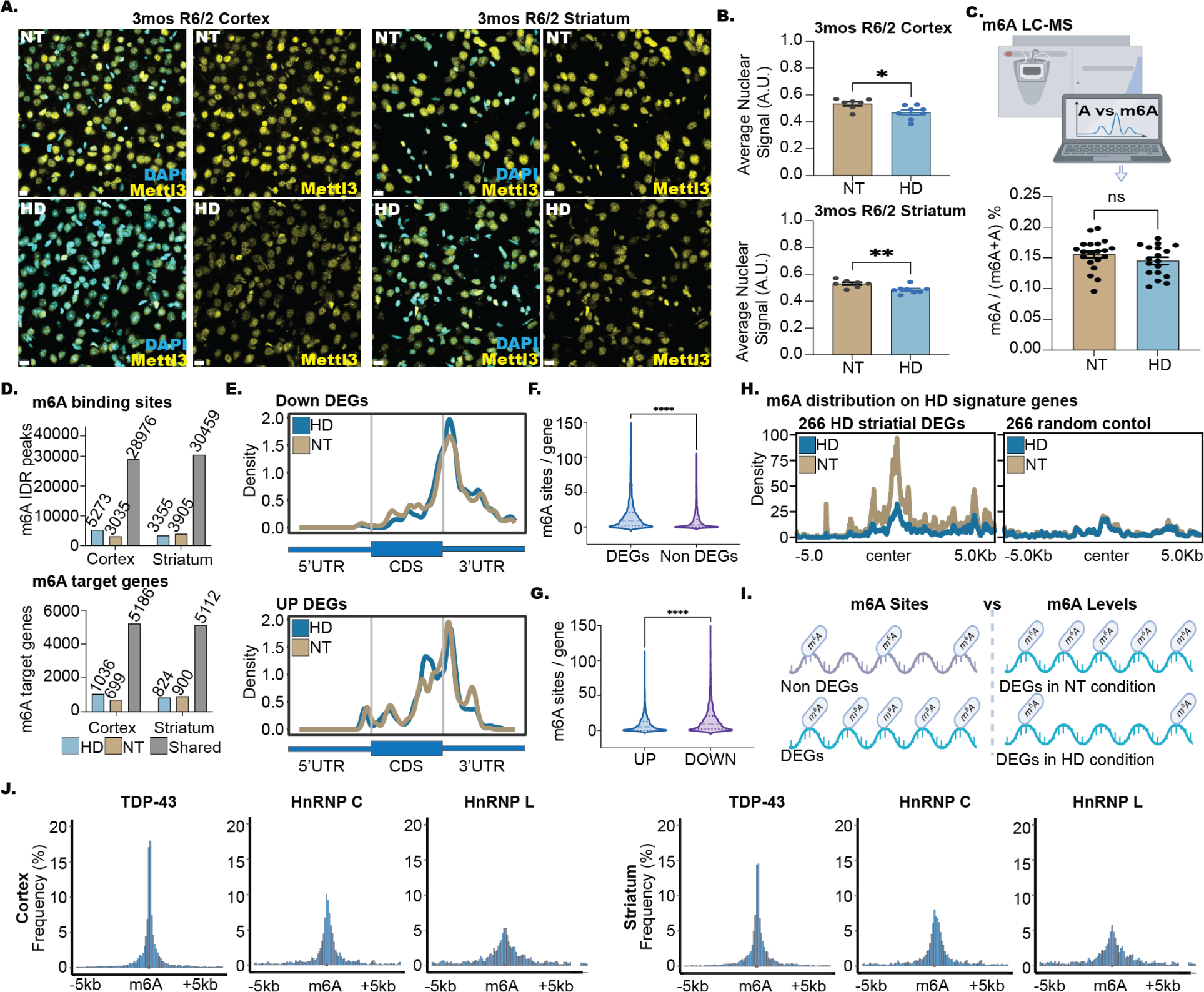
Altered m6A deposition in HD R6/2. (A) Cell profiler analysis of Mettl3 nuclear intensity, statistical significance determined by unpaired t-test, p-value = 0.0184 (cortex), 0.0093 (striatum). n=8 animals per condition were quantified each averaging three cortical or striatal regions. (B) Representative images showing modest but significant (p-value < 0.05) decrease in Mettl3 signal in the HD brains. (C) Mass-spectrometry for m6A shows no significant changes by unpaired t-test (p-value = 0.2025) in global m6A levels. (D) Number of m6A-eCLIP sites detected and genes detected. Numbers ontop of bars represent counts (E) m6A distribution on down and upregulated genes showing altered m6A modifications across transcripts in HD mice. (F) Increased m6A sites per gene on genes that are dysregulated in HD (DEGs), in particular in downregulated genes compared to upregulated genes (G). (H) m6A site deposition (localized levels) on 266 genes dysregulated in HD striatum (I) schematic of how m6A deposition is altered in HD. Unpaired t-test was performed on F&G p-value < 0.0001 (J) Histogram plot showing TDP-43 eCLIP sites relative to an m6A site (from m6A eCLIP data) (0 position) compared to HnRNP C (a known m6A reader protein) and HnRNP L binding motifs. (K) Proposed mechanism of m6A-dependent TDP-43’s binding to HD dysregulated genes.

TDP-43 binding was decreased in genes defining the striatal HD signature. Similarly, m6A deposition decreased in the same striatal genes in the HD condition (Fig. 4H). These important results suggest a direct connection between RNA modification status, TDP-43 binding, and mHTT-dependent transcriptomic alterations; genes that tend to be differentially expressed have more m6A sites, however in HD those same genes are less methylated (Fig. 4I). Given the enrichment of the m6A and TDP-43 binding motifs on HD alternative gene expression changes, we evaluated whether TDP-43 clustered close to m6A sites. Centering the m6A-sites and looking 5 kb upstream and downstream, the TDP-43 binding sites were plotted relative to the m6A-site at the center position. We next looked for known RBP motifs in the genome (HnRNP C – UUUUU / HnRNP L – ACACA) as a comparison and controlled for expressed genes in the R6/2 striatum and cortex (Fig. 4J). TDP-43, compared to HnRNP C and HnRNP L, has a higher center peak overlapping with m6A sites. HnRNP C is a known m6A reader protein that recognizes the m6A motif. TDP-43’s higher peak suggests that TDP-43’s binding may be more dependent on m6A than HnRNP C in HD^75^ and that a decrease in m6A deposition on dysregulated HD genes results in decreased TDP-43 binding and decreased stability. This data reveals a novel mechanistic connection between TDP-43 binding sites and m6A deposition sites in HD, suggesting that m6A methylation may be required for TDP-43 binding.

### Detection of novel unannotated splicing events in the HD R6/2 model

TDP-43 acts as a repressor of cryptic exons (CE) in ALS/FTD with TDP-43 nuclear loss resulting in increased inclusion of CE^34–36,76^. Since we observed TDP-43 colocalization with mHTTex1 nuclear inclusions, reduced nuclear expression of TDP-43 and cytosolic inclusions, we analyzed if CE splicing was affected in HD. We had initially performed rMATs analysis with the novel splicing detection option ^58^. Separating annotated AS events from unknown unannotated events, ∼50% of the AS changes were novel in the R6/2 cortex and striatum (Fig. 5A), not having been previously investigated for these brain regions. We next applied the approach from Brown et al., 2022^35^, and Ma et al., 2022^36^, for novel splicing detection by subjecting our data to MAJIQ and LeafCutter analysis^77,78^. MAJIQ detected 168 (74 NT, 94 HD) novel cortical alternative splicing events, and 211 (113 NT, 98 HD) striatal, after applying the probability cutoff of > 0.90 and P(deltaPSI> 0.1) for discovery (Fig. 5B). LeafCutter detected 164 (73 NT, 91 HD) cortical, and 244 (158 NT, 86 HD) striatal after using an adjusted P-value cutoff of < 0.05 and deltaPSI > 0.1 (Fig. 5C). GO analysis revealed that the genes with novel splicing changes occur in the following categories: modulation of excitatory postsynaptic potential, regulation of cation channel activity, neuromuscular junction development, regulation of dendritic spine morphogenesis, and positive regulation of neuron projection development (Fig. 5D). Using long-read sequencing from the same samples, we found evidence of ∼57% of the novel splice junctions (Fig. 5E). Comparing the novel splicing events from R6/2 to mouse neurons and cardiac cells altered by TDP-43 knockdown from Susnjar et al., 2022^70^, we observed 55% overlap of TDP-43-dependent novel splicing events (192/348) (Fig. 5F). Next, we asked whether these unannotated splicing events could drive differential gene expression changes and indeed, their presence corresponds to gene expression changes in both directions in HD R6/2 (Fig. 5G). To compare altered novel splicing from HD266ssgs with increased and decreased unannotated exon usage in the HD R6/2 mouse data, and corresponding direction of gene expression changes, we use mapped RNA-seq data to generate sashimi plots (Fig. 5H & 5I). When TDP-43 binding sites are overlayed, TDP-43 binding sites are present within the intron containing the unannotated exons, suggesting a potential TDP-43-dependent regulation of expression. Our analysis identified altered TDP-43 localization and novel splicing changes in the HD R6/2 mouse model in genes responsible for neuronal development and function that are dysregulated in HD.

**Figure 5:**
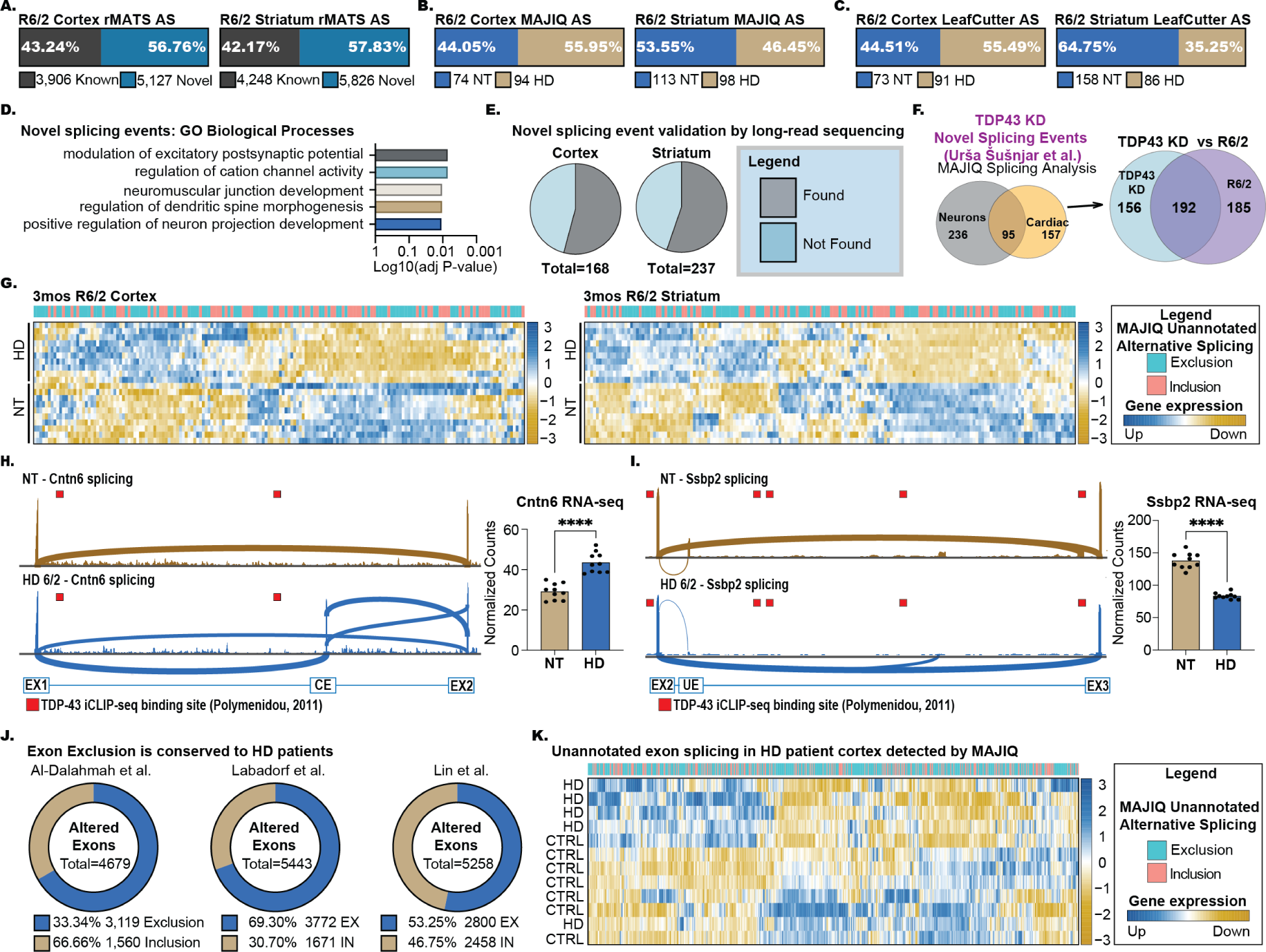
Detection of Cryptic Exons in HD R6/2 and Human RNA-seq data. (A) Bar plot showing rMATS splicing changes that are annotated (known) versus no annotated (novel). (B-C) Bar plots showing MAJIQ and Leafcutter output for the detection of novel unannotated splicing events in the cortex and striatum from 3-month-old R6/2 (NT). (D) Gene Ontology analysis of mHTT dependent novel splicing events for biological processes. (E) PacBio Iso-Seq validation of Novel splicing events, isoforms were searched for the inclusion or exclusion of novel splice products. (F) Venn diagram showing MAJIQ junction overlap between TDP43 KD changes from Susnjar et al. (2022) versus R6/2 cortex and striatum samples. Tables on the right show overlapping genes. (G) Heatmap showing exclusion and inclusion of novel unannotated splicing events in 3-month-old HD R6/2 vs NT (cortex and striatum) and corresponding gene expression changes of the genes that contains the splicing event. (H-I) IGV shashimi plots of increased cryptic exon (CE) splicing in HD (Cntn6, H) or loss of novel unannotated exon (UE) in NT (Ssbp2, I) in HD. (J) Exon skipping molecular signature is preserved in HD patients, across previously published HD RNA-seq datasets ^79–81^. (K) Heatmap showing exclusion and inclusion of novel unannotated splicing events detected by MAJIQ in HD patient cortex compared to non-HD control from Al-Dalahmah et al., 2020. ^80^, and corresponding gene expression changes of the genes that contains the splicing event.

### Altered exon exclusions and novel unannotated splicing events in HD patient brain

With the elucidation of TDP-43 pathology in HD patient brains, we explored if the alternative splicing and altered novel splicing events translated to human subjects. Using three published HD RNA-seq datasets from^79^ (prefrontal cortex Brodmann area 9, Grade III/IV, 20 HD 49 Controls),^80^ (anterior cingulate cortex, grade III/IV, 6 HD, 6 Controls), and^81^ (Broadmann area 4 (BA4) primary motor cortex, grade II-IV, 7HD, 7 Controls), we carried out rMATs & MAJIQ analysis. 12,469 alternative splicing events were detected in BA9 samples, 5,258 in the BA4 samples, and 7,730 in the anterior cingulate cortex samples (FDR < 0.05). We assessed if the splicing signature from mice translated to human subjects and observed higher exon exclusion events detected in the HD condition, confirming that increased exon exclusion is a prominent alternative splicing signature in human HD (Fig. 5J). We next set out to understand whether novel unannotated splicing events are altered in human HD. Through MAJIQ analysis, we detected 78 events (42 Controls, 36 HD) in the prefrontal cortex and 118 events (56 Controls, 62 HD) in the cingulate cortex. Similar to the HD R6/2, the novel unannotated splicing changes in the human data contribute to differential gene expression changes in both directions (Fig. 5K). This human HD RNA-seq analysis thus confirms the mouse finding of increased exon exclusion events and altered novel unannotated splicing in HD.

## Discussion

The dysregulation of TDP-43 in ALS and FTLD is an active area of investigation with development of biomarkers and therapeutic strategies such as ASOs to restore appropriate splicing to TDP-43 targets^82^. While potential splicing changes have been documented in HD^81,83^, including the incomplete splicing of HTT RNA to expression Httex1a protein, the underlying mechanisms and the potential involvement of TDP-43 as a potential driver of mis-splicing have not been defined. Here we show that TDP-43 loss of function together with aberrant m6A modification may be a driver of mis-splicing in HD, leading to the altered expression of critical striatal genes dysregulated in HD. Building on our previous data showing altered TDP-43 localization in human patient brains ^18^, we generated and integrated novel multi-omics data herein, including RNA-sequencing, RASL-seq, Pacbio long-read sequencing, m6A-eCLIP sequencing, and TDP-43 eCLIP-sequencing to investigate mechanisms involved in aberrant splicing. We demonstrated that the RBP TDP-43 and the m6A writer, METTL3, have altered protein subcellular localization and protein expression, respectively, in R6/2 mice. These alterations accompanied a corresponding enrichment in HD-specific alternative splicing and decreased interaction with dysregulated striatal HD signature RNAs. Immunofluorescent (IF) imaging in HD mice and HD patient brain tissue revealed co-localization of TDP-43 with mutant HTT nuclear inclusions and decreased nuclear TDP-43 and corresponding increase in aggregated phosphorylated TDP-43 in the cytoplasm. Our analysis of RNA-seq data from both mouse and human samples revealed changes in alternative splicing with increased exon exclusion events in the HD condition.

We observe that 60% of the altered splicing events occur within genes that are differentially expressed in HD with the majority leading to decreased gene expression. We hypothesize that mHTT can disrupt HTT’s normal interactions with RBPs, including TDP-43 and METTL3, that may result in regulatory RBPs not being properly bound to their RNA targets, thus altering splicing and leading to dysregulated AS observed in our study. Consistent with this hypothesis, we show that TDP-43 binding to genes crucial to striatal neuronal maturation and survival is decreased in the HD condition, suggesting that TDP-43’s binding provides stability to target transcripts. m6A-seq revealed increased m6A modifications found on genes differentially downregulated in HD compared to non-differentially expressed genes. However, m6A depositions at specific sites on these genes in HD are decreased. This finding synergizes with the TDP-43 CLIP-seq data, as we report that TDP-43 binding is enriched near m6A modification. A functional crosstalk between RBPs and m6A has been shown for Tau, where HNRNPA2B1 connects oligomeric Tau with m6A modification as a linker^84^. Further, m6A modification is required for TDP-43 binding to RNA in ALS spinal cord^52^ and toxicity modulated through genetic perturbation of m6A machinery. Our finding that TDP-43 binding corresponds with m6A deposition on downregulated striatal genes in HD suggests a novel co-regulatory role for m6A modification with TDP-43 in HD.

We also report expression of novel unannotated exon splicing in both HD mouse and human brain RNA, consistent with disruption of TDP-43s known function as a negative regulator of cryptic exons^85–87^. Typically, the nuclear translocation of TDP-43 and aggregation of phosphorylated TDP-43 in the cytoplasm causes an increase in CE regulation of in STMN2 and UNC13a^35,36,38,82,87^. Using isogenic iPSC-derived striatal enriched neurons, we detected altered novel exon usage in the HD condition that was similar to the effect of TDP-43 KD in control iPSC-neurons. Further, there was a significant overlap with previously annotated TDP-43 KD studies in mice^23^. The presence of aberrant novel unannotated exon splicing corresponds to hallmark HD genes that are primarily downregulated, however, unexpectedly, both an increase and decrease in novel unannotated exon expression were identified to result in differential gene expression. This proposes a mechanism in which HTT’s interaction with multiple RBPs regulates CE splicing. Further studies are required to identify additional CE-regulating RBPs.

The detection of nuclear pTDP-43 AL-bodies in our study may provide the field with a TDP-43 pathology potentially unique to HD. These circular aggregation-like bodies do not morphologically resemble known normal nuclear aggregates^88^. The co-localization of AL-bodies with HTT only in neurons lacking canonical HTT nuclear inclusions suggests the possibility that aggregation of TDP-43 in HD is dependent on mHTT, until mHTT reaches a seeding density to form the conical nuclear inclusions. When this happens, we hypothesize that TDP-43 may then translocate into cytoplasmic aggregates. The shape of AL-bodies suggests an unidentified core, similar to PML bodies, forming a donut-like structure around ubiquitinated and sumoylated proteins^89^. Alternatively, AL-bodies may co-localize with the nucleolus (89). Finally, AL bodies may represent RNA-free TDP-43-containing anisosomes within the nucleus that we have previously shown can occur via proteosome inhibition or specific mutation in TDP-43’s RRM^90^. Our TDP-43 eCLIP-seq detected decreased TDP-43 binding to target RNAs, making it plausible that these AL-bodies may contain RNA-free TDP-43. Here, we identified three TDP-43 aggregation phenotypes in HD, one of which has not previously been observed, which presents the possibility that our molecular readouts are resulting from a combinatorial effect of both known and novel TDP-43-dependent regulation. Our future efforts will be aimed at elucidating the role of AL-bodies role to HD progression and neurodegeneration.

In summary, we present a systematic analysis of mechanisms that may contribute to aberrant splicing in HD, including through post-transcriptional RNA processing (see schematic, Fig. 6). The data supports TDP-43 loss of function coupled with altered m6A modification as a novel mechanism underlying alternative splicing/cryptic exon usage and critical gene dysregulation implicated in HD. Neuropathologically, we show a novel aggregate-like nuclear body in human HD that is independent of classical nuclear inclusions. Finally, the body of work suggests that the impact of TDP-43 dysregulation involved as a driver of disease pathogenesis is broad and may represent a pivotal therapeutic target for neurodegenerative diseases in general. The herein discovered relationship between TDP-43 and m6A may also suggest emergent methods to control m6A could have a therapeutic benefit through regulation of TDP-43^91–93^.

**Figure 6:**
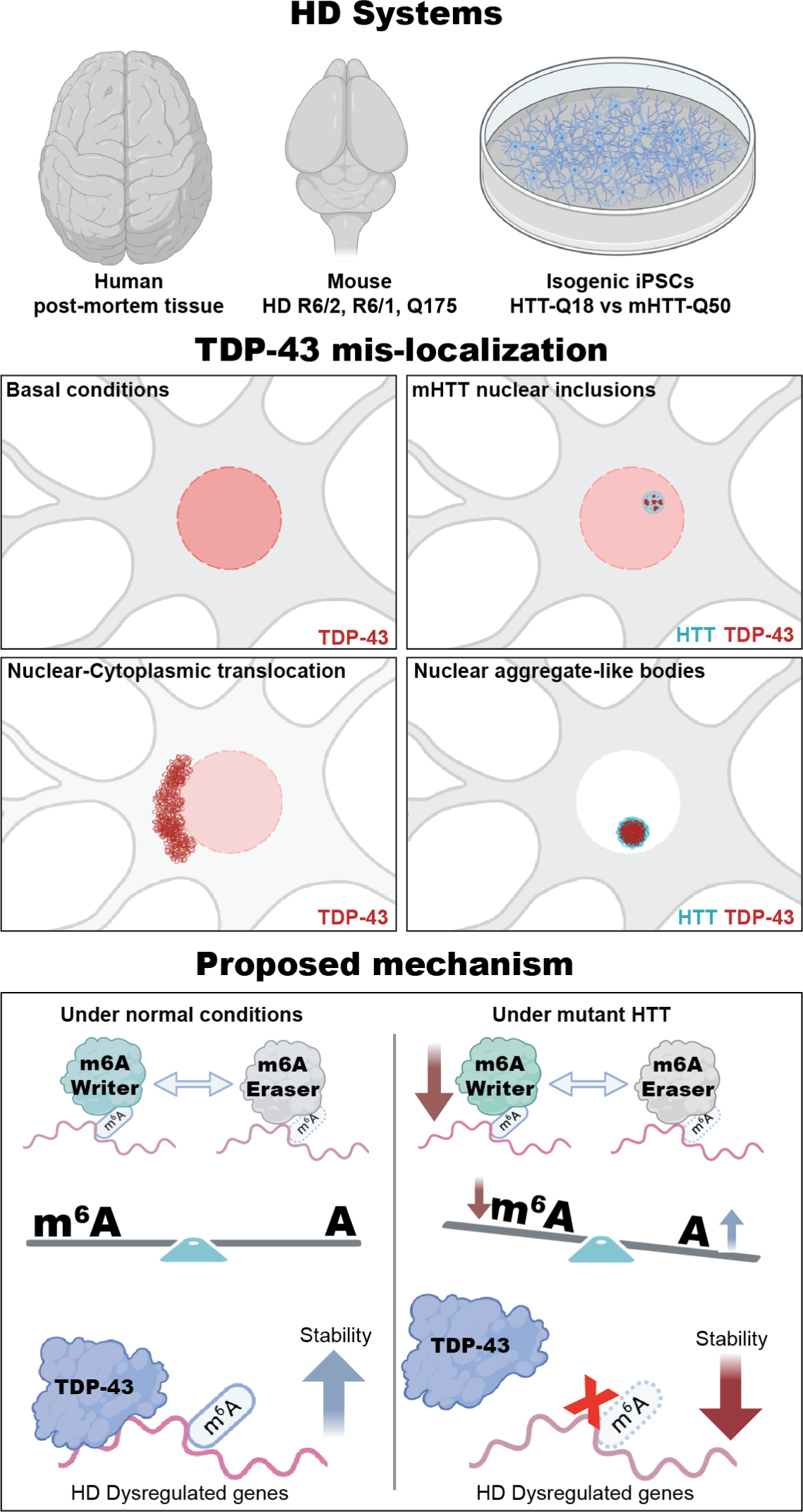
Summary of study findings and proposed mechanisms. Summary Schematic showing the model systems used in this study, phenotypes, and proposed mechanisms. We show four different TDP-43 conditions - basal nuclear localization in unaffected brain, colocalization with classic mHTT nuclear inclusions, mislocalization of TDP-43 to the cytoplasm and pTDP-43 cytosolic aggregates and a novel nuclear aggregate-like body (AL body) in the nucleus comprised of HTT and TDP-43. We hypothesize that the dysregulation and imbalance of TDP-43 and m6A results in altered splicing and stability of signature striatal genes in the HD condition.

## Materials and Methods

### Animal model, handling, and tissue harvest

Experiments were carried out in accordance with the ‘Guide for the Care and Use of Laboratory Animals of the NIH’ and an approved animal research protocol by the ‘Institutional Animal Care and Use Committee (IACUC)’ at the University of California, Irvine an AAALAC accredited institution. R6/2 and non-transgenic control male and female? mice were obtained from Jackson Laboratory at ∼5 weeks of age. All mice were housed on a 12/12-hr light/dark schedule with *ad libitum* access to food and water. Animals were aged then euthanized with Euthasol overdose (pentobarbital sodium and phenytoin sodium). Cardiac perfusion was performed with 0.01 M PBS, followed by brain harvesting and isolation of Striatum and Cortex from the left hemisphere that was flash-frozen in liquid nitrogen and stored at -80°C until use for biochemical analysis. The other halves were post-fixed in 4% paraformaldehyde, cryoprotected in 30% sucrose, and cut at 30 μm on a sliding vibratome for immunohistochemistry (IHC) described below. For Biochemistry frozen tissues were lysed (Lysis Buffer: 50 mM Tris-HCl pH 7.4,100 mM NaCl, 1% NP-40 (Igepal CA630), 0.1% SDS, 0.5% sodium deoxycholate, 1:200 Protease Inhibitor Cocktail III (add fresh), 0.1 mM PMSF, 25 mM NEM, 1.5 mM aprotinin, and 23.4 mM leupeptin). In brief samples were homogenized by douncing in lysis buffer followed by incubation on ice for 30 minutes. Lysate were then sonicated 3X 10seconds @40% amplitude. Protein quantification was performed by Lowery protein assay with linear range dilution.

### Differentiation of iPSCs to medium spiny neurons and siRNA knockdown of TDP-43

iPSCs were derived from fibroblasts from a non-affected patient using nonintegrating reprogramming techniques and CRISPR modified to an isogenic series with 18Q and 50Q CAG repeat length. Cells are grown on Matrigel coated plates in tissue culture sterile conditions and passaged with nonenzymatic dissociation medium. Cells are fed with mTeSR1 (Stem cell technologies) every day, and passage at ∼70% confluency. Small molecule-based Differentiation to MSNs were followed by Smith-Geater et al., 2020 and outline in (Fig. 3B). Human TDP-43 and non-targeting control siRNA were obtained from (Accell SMARTPool, Horizondiscovery, cat# E-012394-00-0050), cells were treated with siRNA on day 23 and harvested at day 37.

### Co-Immunoprecipitation / immuno-blotting

For each tissue type the optimal protein concentration and primary antibody concentration were determined by linear range according to Li-COR’s protocol. For Co-IP 1mg of protein was used for each IP, 1:1000 dilution of primary antibody, 30 ul of Dynabeads sheep anti-rabbit or Goat anti-mouse. IP was carried out by incubation for 1hr at room temp then a wash performed with 3X high salt wash buffer (50 mM Tris-HCl pH 7.4, 1 M NaCl, 1 mM EDTA, 1% NP-40, 0.1% SDS, 0.5% sodium deoxycholate), followed by 2X low salt wash buffer (20 mM Tris-HCl pH 7.4, 10 mM MgCl2, 0.2% Tween-20). Elution was achieved by 10-minute incubation at 80°C in 1X LDS and 1mM DTT. Co-IP and regular western blot samples were run on 4-12% Bis-Tris gel and 3-8% tris-acetate. Specific protein band analysis was performed using the Li-COR EmpiraStudio software with normalization to REVERT total protein stain.

### Immunofluorescence staining

A list of primary and secondary antibodies with dilutions used in this study can be found in supplemental table S9.

For mouse samples coronal sections that included the striatum were selected, Antigen Retrieval was performed for 20minutes at 80C (AR buffer: 10 mM Tri-Na Citrate buffer, pH9 + Tween20 0.05%). Tissue slices were permeabilized for 10 minutes @RT (Permeabilization buffer: PBS + 2.5% BSA. _ 0.2% Triton X-100), followed by blocking for 2 hours @RT (blocking buffer: PBS + 5% NGS (or NDS) + 1% BSA + 0.1% Triton X-100). Primary antibodies were added at the indicated concentration in blocking buffer and incubated overnight @4°C. Secondary antibody was performed for 2 hours @RT followed by Hoeschst (1:3000) for 10 minutes @RT. Tissues were then mounted onto slides and coverslips with Fluoromount-G (southern biotech cat# 0100-01) and stored at 4°C. For human, 5microns Paraffin-embedded sections were used. Tissue sections were heated at 65C for 30min then deparaffinized with 100% CitriSolv (Fisher Scientific, 04-355-121) for 15mins X2, 100% ETOH for 5mins X2, 95% ETOH for 5mins, 70% ETOH for 5mins, 50% ETOH for 5mins, miliQ H20 for 5 mins X 2, rehydrated, and antigen retrieval was performed with Vector Laboratories cat.no. H-3301) 20 minutes at 95°C. Sections were blocked for 1 hour @RT with 5% Normal Goat or Donkey Serum in 0.1% Triton X-100. Sections were incubated in primary antibody overnight at 4°C in 1% Normal Donkey Serum in 0.1% Triton X-100. Sections were then incubated in secondary antibodies (1:400 dilution) for 1 hour @RT in 1X PBS. Secondary antibodies were used: Alexa Fluor 488 (1:400, cat. #A-21202, ThermoFisher), Alexa Fluor 555 (1:400, cat. #A-31570, ThermoFisher). Tissues were then treated TrueBlack Lipofuscin Autofluorescence Quencher (cat. #23007, Biotium), and incubated in Hoeschst for 10 minutes @RT. Sections were then mounted with coverslips were mounted using Fluoromount-G.

### Microscopy, IF intensity measurement, and Co-localization analysis

Images were taken on the Zeiss AiryScan 900 and Olympus Fluoview FV2000 confocal system, with the 40X and 63X objective, images were processed using the AiryScan software. Images were taken with the same acquisition settings. Images were then imported to Imaris imaging software version 9 for post-imaging analysis. For image analysis, images had all ‘auto-adjustment’ settings reset to raw values. Next, the exact brightness and contrast, min/max, and gamma (default value of 1) were applied to all images for comparison and analysis. For Intensity measurement, each cell containing TDP43 nuclear signal was quantified with the Imaris surface tool. Normalization was performed between animals by dividing by surface volume. Statistical analysis was performed with unpaired t-test between HD vs NT.

### RNA library preparation & Sequencing

Starting from mouse frozen cortex, striatal tissue, or iPSCs, 1ug of RNA was extracted and DNase treated using Trizol (Invitogen) and the manufacturers procedure, the isolated RNA was then used for paired-end library preparation with Illumina Truseq with Ribo-Zero and sequenced on the Illumina Novaseq-6000 using the S4 flowcell. RNA quality and concentration was determined with Bioanalyzer and quantified with Qubit.Targeted read depth was 50 million reads per samples using paired-end sequencing 100bp. fastq files were subjected to FASTQC for QC analysis, aligned to mm10 or hg38 using STAR aligner (v2.7.0). Count matrix was generated using featurecounts and differential gene analysis was performed with DESeq2 and genes with low expression were filtered out. For plotting of normalized counts, counts were normalized with Trimmed Mean of M-values (TMM) in the edgeR package.

### Alternative splicing, & Cryptic exon analysis

rMATS was used for analysis of alternative splice sites from RNA-seq data. The following options were used [“ --od “nt_vs_hd” --tmp “tmpFolder” -t paired --readLength 101 --cstat 0.0001 --libType fr-firststrand --nthread 20 –novelSS]. For novel splice sites rMATS was reran with the option – NovelSS. For analysis of alternative splicing of single nuclei RNA-seq data, DESJ-detection was used according to S. Liu et al., 2021. In every instance, only splice sites with a False Discovery Rate < 0.05 were considered significant. MAJIQ and Leafcutter analysis were performed following A.L. Brown et al., 2022 ^35^ and X.R. Ma, et al., 2022 ^36^.

### RASL Sequencing

RASL-seq was carried out as outlined by H Li et al., 2012^94^. In brief HD R6/2 (n=3 NT, 3 HD), the Q150 (n=5 WT, 4 HET, 3 HOMO)^61^, and Q175 (n=5 WT, 4 HET, 4 HOMO)^61^ mouse models at 3, 7, and 18 months, respectively were used to construct libraries. 9,496 primer pairs targeting known splice junctions were used to amplify libraries for illumina sequencing platform. Standard analyses were performed as outline by H Li et al., 2012^94^.

### PacBio Iso-Seq

10ug of Trizol-extracted RNA from mouse cortex and striatum was treated with Terminator 5’-Phosphate dependent exonuclease (Lucigen TER51020) according to manufacturer’s protocol., RNA was purified by Zymo RNA Clean and Concentrator-5. Reference transcripts SIRV set 3 (Lexogen) diluted 1:10 and 0.3 ul was added to 300 ng of purified RNA prior to cDNA synthesis. A modified protocol was used for cDNA synthesis using superscript IV and template switching oligos ^95^. cDNA was pre-amplified using KAPA HiFi HotStart Ready mix (Roche) for 27 cycles. Iso-seq libraries were prepares using the SMRTbell Template Express Prep Kit 2.0 (Pacific Biosciences) using 300 ng cDNA input per sample according to the manufacturer’s protocol. Following library preparation, DNA is purified with 0.45x AMPure PB beads, washed with 80% ethanol twice and eluted with 100ul of elution buffer. DNA was eluted from the beads at 37°C for 15 minutes. A second round of purification with AMPure PB beads was included and each DNA library was eluted in 10ul. DNA library quality and consistency were verified with Qubit and Bioanalyzer and loaded onto the PacBio Sequel II sequencer. PacBio sequencing data was analyzed using SMRT Analysis software (Pacific Biosciences). Consensus reads were generated using ccs 4.2.0 with options --skip-polish --minLength 10 --minPasses 3 --min-rq 0.9 --min-snr 2.5. Adapters were removed with lima 1.11.0 with options --isoseq --min-score 0 --min-end-score 0 --min-signal-increase 10 --min-score-lead 0. Full-length non-chimeric reads were extracted, were oriented to the correct strand, and had poly-A tails clipped using isoseq3 refine 3.3.0 with options --min-polya-length 20 --require-polya. The resulting full-length non-chimeric reads were mapped to mm10 reference genome using minimap2 version 2.17-r941 (doi:10.1093/bioinformatics/bty191) with options -ax splice:hq -uf -MD. TranscriptClean v2.0.2 (doi.org/10.1101/672931) was used for error correction with option --canonOnly. and TALON v5.0 (doi.org/10.1101/672931) was used for error correction and transcript identification and quantification. The module talon_label_reads with option --ar 20 was used to compute the fraction of As at the ends of read alignments. TALON databases of mouse (Ensembl 87 annotations) were created using talon_initialize_database with options --l 0 --5p 500 --3p 300. The talon module was run with default parameters to identify transcripts using the initialized database. Filtered transcript lists, which are limited to known and consistently observed transcripts, were generated using talon_filter_transcripts. Filtered and unfiltered transcript abundances were obtained using the talon_abundance module.

### TDP-43 and m6A eCLIP-seq

Males and Females R6/2 and corresponding non-transgenic animals from matching cohort (n=4 per group) were aged to 12 weeks of age, Striatum and Cortex were dissected and flash frozen in liquid nitrogen. For m6A-eCLIP, total RNA was extracted using Trizol, mRNA was enriched using poly-dT beads (NEB) and cleaned with Zymo clean and concentrator 5 according to standard protocol, then followed ECLIPSE BIO protocol. For TDP43-eCLIP samples were homogenized to a powder with BioSpec 59012N Tissue Pulverizer. Samples were next UV crosslinked 2X @400mj/cm^2^. Samples were collected in 1.5mL tube. Next the RBP-eCLIP kit from ECLIPSE BIO with data analysis was used following the RBP-eCLIP Protocol V2.01R to generate TDP-43 eCLIP sequencing libraries (based on Van Nostrand et al., 2016 Nature Methods). 2ug of TDP-43 Antibody was used for each IP (Bethyl A303-223A). IP and corresponding input samples were sequenced at 50 million reads using 100BP paired-end sequencing on the Novaseq-6000 with S4 flow cell. Male and female samples were compared by using Irreproducibility Discovery Rate (IDR) incorporated into CLIPPer peak finder^96^ to identify reproducible binding peaks and filtered by p-value < 0.001 and log(foldchange) > 3. Metagene plot was created with MetaPlotR using the following standard pipeline^97^.

### LC-MS analysis of m6A

Processing of RNA for LC-MS-based m6A analysis was performed as described in Mathur et al. (PMID: 34401789). Briefly, 100 ng of twice-purified poly(A)-RNA was digested with 1 unit of nuclease P1 (Cat# N8630, Sigma-Aldrich) for 2 h at 37 °C, followed by treatment of 1 unit of alkaline phosphatase (Cat# P5931, Sigma-Aldrich) for 2 h at 37 °C. 21.5 μL of the purified nucleoside sample (equivalent of 25 ng RNA) was mixed with 2X volume (43 μL) of acetonitrile. Samples were centrifugated at 16,000g for 10 min at 4 °C, and 40 µL of supernatant was loaded into MS vials. Adenosine and m6A signals were analyzed by quadrupole-orbitrap mass spectrometer (Thermo Fisher Scientific) coupled to hydrophilic interaction chromatography (HILIC) via electrospray ionization. LC separation was on an Xbridge BEH amide column (2.1 mm x 150 mm, 2.5 µm particle size, 130 Å pore size; Waters) at 25°C using a gradient of solvent A (5% acetonitrile in water with 20 mM ammonium acetate and 20 mM ammonium hydroxide) and solvent B (100% acetonitrile). The flow rate was 200 µL/min. The LC gradient was: 0 min, 75% B; 3 min, 75% B; 4 min, 50% B; 7 min, 50% B; 7.5 min, 75% B; 11min, 75%. Autosampler temperature was set at 4°C and the injection volume of the sample was 3 μL. MS data were acquired in a positive ion mode with a full-scan mode from m/z 240 to 290 with 140,000 resolution. Data were analyzed using El-MAVEN software (version 0.12.0). Adenosine (Cat# A9251, Sigma-Aldrich) and m6A (Cat# S3190, Selleckchem) standards were quantified based on standard calibration curves using authentic standards.

## Acknowledgments

We would like to thank the families and patients of those affected by HD for their critical contributions to research. We would like to thank the Netherlands Brain Bank and Dr. Richard Faull (New Zealand Brain Bank) for supplying the human brain tissue for our study. Thank you to Dr. Len Petrucelli for the generous gift of the S409/S410 TDP-43 antibody. We would like to thank Anna-Leigh Brown for the help on MAJIQ analysis and scientific discussions. We would like to thank the lab of Dr. Gene Yeo and ECLIPSE bioinnovations for the wonderful scientific discussions.

## Funding

This work was supported by a Chan Zuckerberg Initiative Collaborative Pairs grant (LMT and RS) as well as supported by the following NIH grants: R35 NS116872 (LMT), R01 NS112503 (DWC and CLT), R01 NS124203 (CLT and DWC), R01 NS27036 (DWC), R01 AA029124 (CJ), NIH K22CA234399 (GL), and DOD grant TS200022 (GL). Additional support was provided from the National Institute of Neurological Disorders and Stroke of the NIH under Award Number F31NS124293T32 (TBN), Hereditary Disease Foundation postdoctoral fellowship (RM2), and postdoctoral fellowship from the ALS Association (SVS). The content is solely the responsibility of the authors and does not necessarily represent the official views of the National Institutes of Health.

## Author contributions

Thai B. Nguyen (TBN), Ricardo Miramontes (RM1) Carlos Chillon-Marinas (CCM), Roy Maimon (RM2), Sonia Vazquez Sanchez (SVS), Alice L. Lau (ALL), Nicolette R. McClure (NRM), Whitney E. England (WEE), Monika Singha (MS), Ki-Hong Jang (KJ), Sunhee Jung (SJ), Jharrayne I. McKnight (JIM), Leanne N. Ho (LNH), Maurice A. Curtis (MAC), Richard L.M. Faull (RLMF), Joan Steffan (JS), Jack C. Reidling (JCR), Cholsoon Jang (CJ), Gina Lee (GL), Don W. Cleveland (DWC), Clotilde Lagier-Tourenne (CLT), Robert C. Spitale (RCS), and Leslie M. Thompson (LMT)

Conceptualization: TBN, CCM, RM2, SVS, JCR, DWC, CLT, RCS, LMT

Methodology: TBN, RM1, CCM, RM2, SVS, WEE, MS, KJ, SJ, CJ, GL

Investigation: TBN, RM1, CCM, RM2, SVS, ALL, NRM, WEE, MS, KJ, SJ, JIM, LNH

Visualization: TBM, RM, CCM, RM1, SVS, CLT

Funding acquisition: CJ, GL, DWC, CLT, RCS, LMT

Project administration: TBN, JCR, CJ, GL, DWC, CLT, RCS, LMT

Supervision: JCR, CJ, GL, DWC, CLT, RCS, LMT

Writing – original draft: TBN, CCM, SVS, JCR, RCS, DWC, CLT, LMT

Writing – review & editing: TBN, CCM, SVS, JCR, CJ, GL, RCS, DWC, CLT, LMT

## Competing interests

Authors declare that they have no competing interests

## Data and materials availability

All data are available in the main text or the supplementary materials. All RNA sequencing data will be available in GEO.

## Supplementary Materials

**1. Supp_Tables.xlsx**

## Notes

### Competing Interest Statement

The authors have declared no competing interest.

