## Supplemental Figures for "Aberrant splicing in Huntington’s disease via disrupted TDP-43 activity accompanied by altered m6A RNA modification"

### Supplemental Figure 1: Identification of expression changes in striatum and cortex from HD R6/2 mice

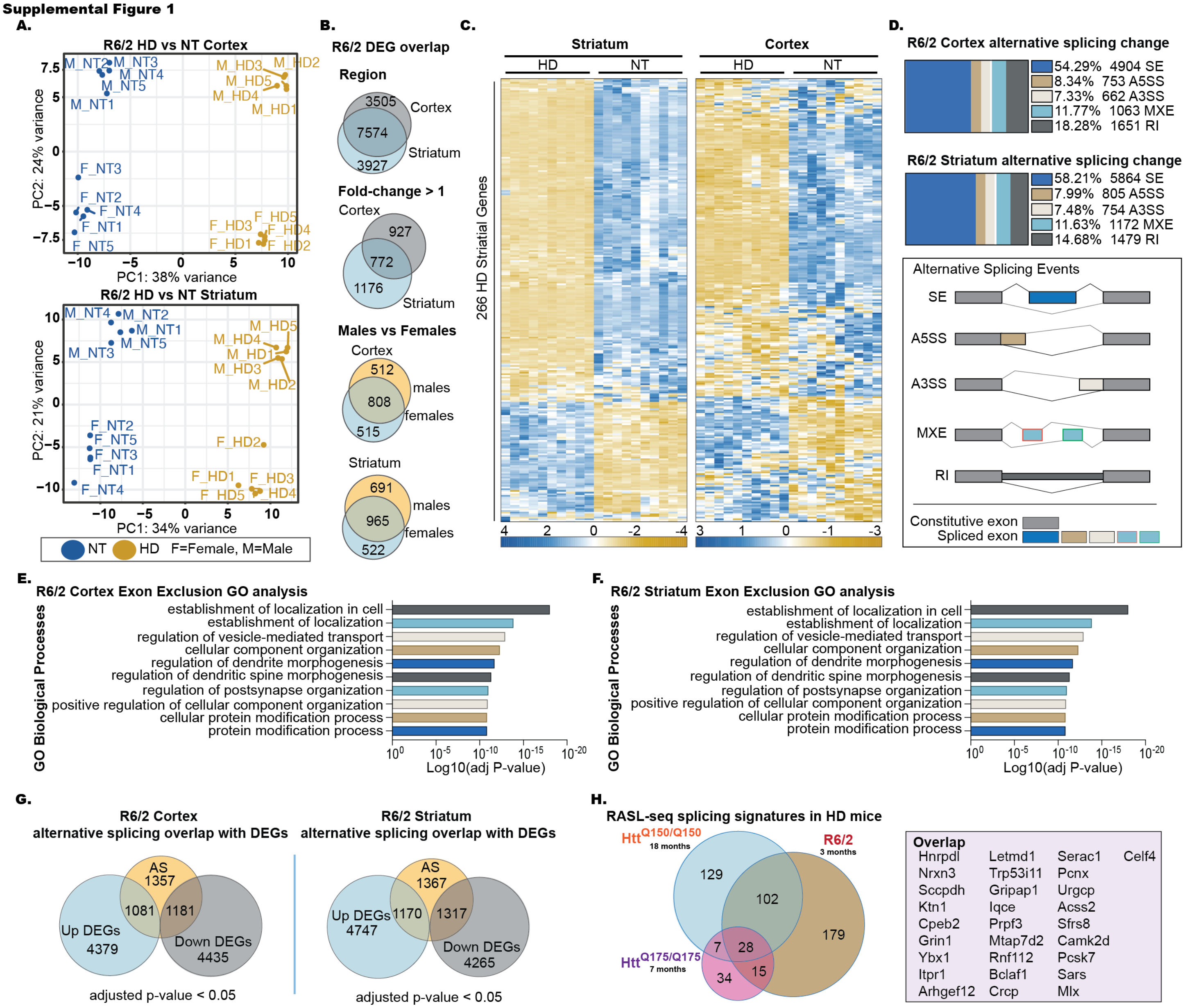


Supplemental Figure 1: (A) Principal Component Analysis (PCA) from the RNA-seq data from striatum and cortex of 12 weeks old R6/2 mice. Separation between groups is observed on PC1 by genotype and on PC2 by sex. Count data was generated with FeatureCount, blue are NT samples and yellow are HD samples. F = female / M = male. (B) Venn diagram showing overlap of differentially expressed genes (DEGs) between striatum, cortex, male, and female samples. Significance was determined by adjusted p-value < 0.05, male vs female DEGs are log2foldchange > 1. (C) Heatmap showing gene expression changes in RNA-seq data of the 266 striatal DEGs defining the signature reported by Obenauer et al. (*46*). Heatmap contains 10 males and 10 female samples with 5 NT and 5 HD from each group. (D) Significant splicing event changes in the cortex (top) and striatum (bottom) from rMATs analysis (FDR < 0.05). Gene Ontology analysis for biological process of the significant skipped exons in the R6/2 cortex (E) and striatum (F). Y-axis displays GO terms, X-Axis represents the -log10(p-value). (G) Overlap between significant alternatively splicing events from rMATs and genes up and downregulated in the R6/2 cortex (left) and striatum (right). (H) Venn diagram showing the overlap of significant splicing changes identified by RASL-seq between the R6/2, Q150 and Q175 mice.

#### Supplemental Figure 2: Decreased TDP-43 nuclear signal in HD

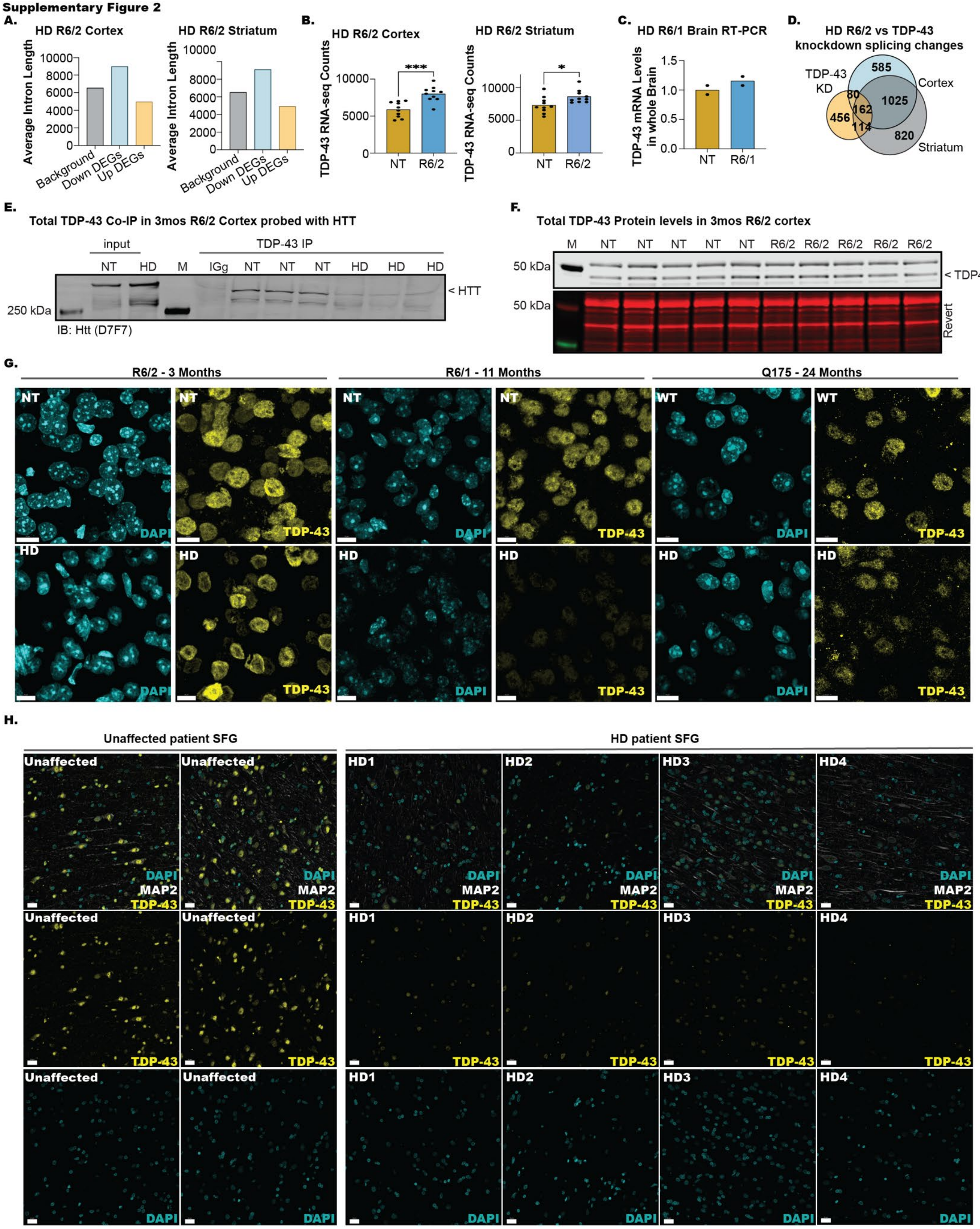

Supplemental Figure 2: (A) For all expressed genes (base mean > 1, from DeSeq2) the intron length was pulled using the mm10 mouse genome annotation and the average intron length per group was calculated (all introns / sum of introns). Intron length in genes differentially expressed in HD mice compared to controls (Up DEGs and Down DEGs) was compared to intron length in non-differentially expressed genes (Background). (B) RNA-seq counts for TDP-43 mRNA in cortex from 12-week-old R6/2 mice generated from FeatureCount. (cortex DESeq2: log2foldchange = 0.25, adjusted p-value 1.66E^-8. Striatum DESeq2: log2foldchange = 0.20, 8.40E^-13) Significance determined by unpaired t-test p-value = 0.0003 (C) RT-PCR of TDP-43 mRNA in whole brain from 11-month-old R6/1 mice and non-transgenic (NT) controls. (D) Comparison of HD rMATs SE output and TDP-43-dependent rMATs SE output (Šušnjar, 2022) in mouse neurons. (E) Western blot by Li-COR using 5ug of lysate per lane from cortex of 3-month-old NT and HD R6/2 males. Red staining shows REVERT total protein staining for loading control. Arrow indicates expected TDP-43 band. (F) Co-immunoprecipitation using a TDP-43 antibody followed by Western blot analysis with an HTT antibody using lysates from age matched (3-month-old) NT control or HD R6/2 animals. (G) Immunofluorescence (IF) staining of the cortex of 3-month-old HD R6/2 mice (left), 11-month-old HD R6/1 mice (middle), 24-month-old Homozygous Q175 mice (right) and wild-type littermates (NT or WT) scalebar=10um. Cyan is the nuclei marker DAPI, yellow shows total TDP-43 staining. (H) Representative IF staining images of superior frontal gyrus from HD patients compared to non-HD control individuals showing decreased TDP-43 (yellow) signal intensity. scalebar=20um.

**Supplemental Figure 3: Accumulation of phosphorylated aggregated TDP-43 and TDP-43 co-localization with HTT nuclear inclusions.**

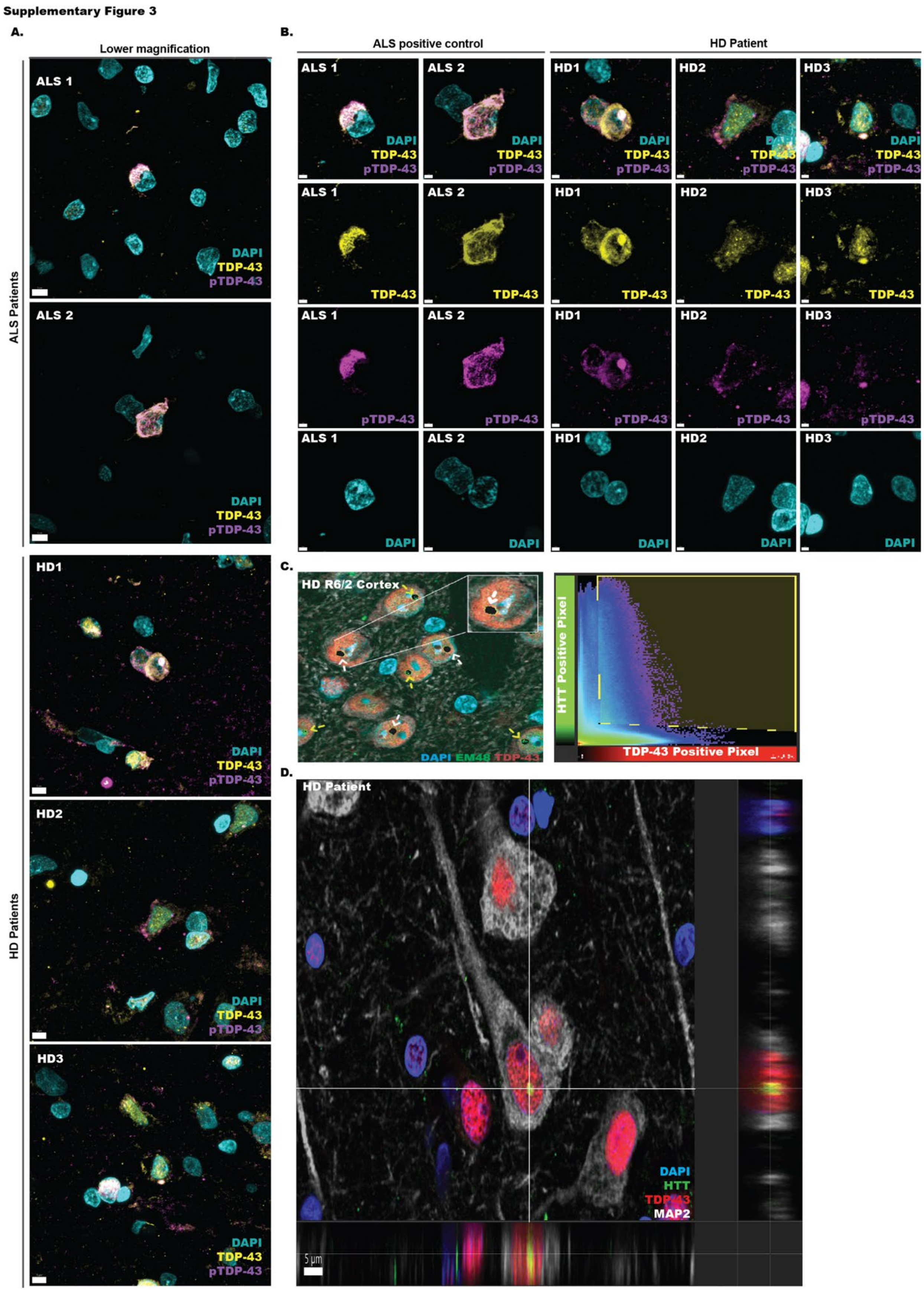

Supplemental Figure 3: (A) Representative low magnification IF staining images in the motor cortex from an ALS patient (positive control) compared to the superior frontal gyrus (SFG) of HD patients, using antibodies against total TDP-43 (yellow), phosphorylated TDP-43 (purple), and nuclear stain DAPI. scalebar=5um (B) Representative high magnification IF staining images in the motor cortex from an ALS patient (positive control) compared to the superior frontal gyrus (SFG) of HD patients, using antibodies against total TDP-43 (yellow), phosphorylated TDP-43 (purple), and nuclear stain DAPI. scalebar=2um (C) Representative images that underwent IMARIS co-localization analysis, left images are from R6/2 cortex, black dots are areas in which TDP-43 and EM48 (HTT) can be detected at the pixel levels. Right graph shows the amount of TDP-43 and EM48 (HTT) signal from each pixel within the black area from the left image. Yellow highlighted area captures pixels that contains signal from both stains. Analysis was done in a 2D slice from a 3D image. The white arrows represent complete co-localization, the yellow arrows represent partial co-localization, black masks reflect pixels selected on the histogram. (D) Confocal imaging of superior frontal gyrus (SFG) from HD patients showing modest colocalization of TDP-43 (red) with HTT nuclear inclusions (green) detected with an N-terminus HTT antibody within MAP2 (white). scalebar=5um

**Supplemental Figure 4: Aggregation-Like phosphorylated TDP-43 nuclear Bodies**

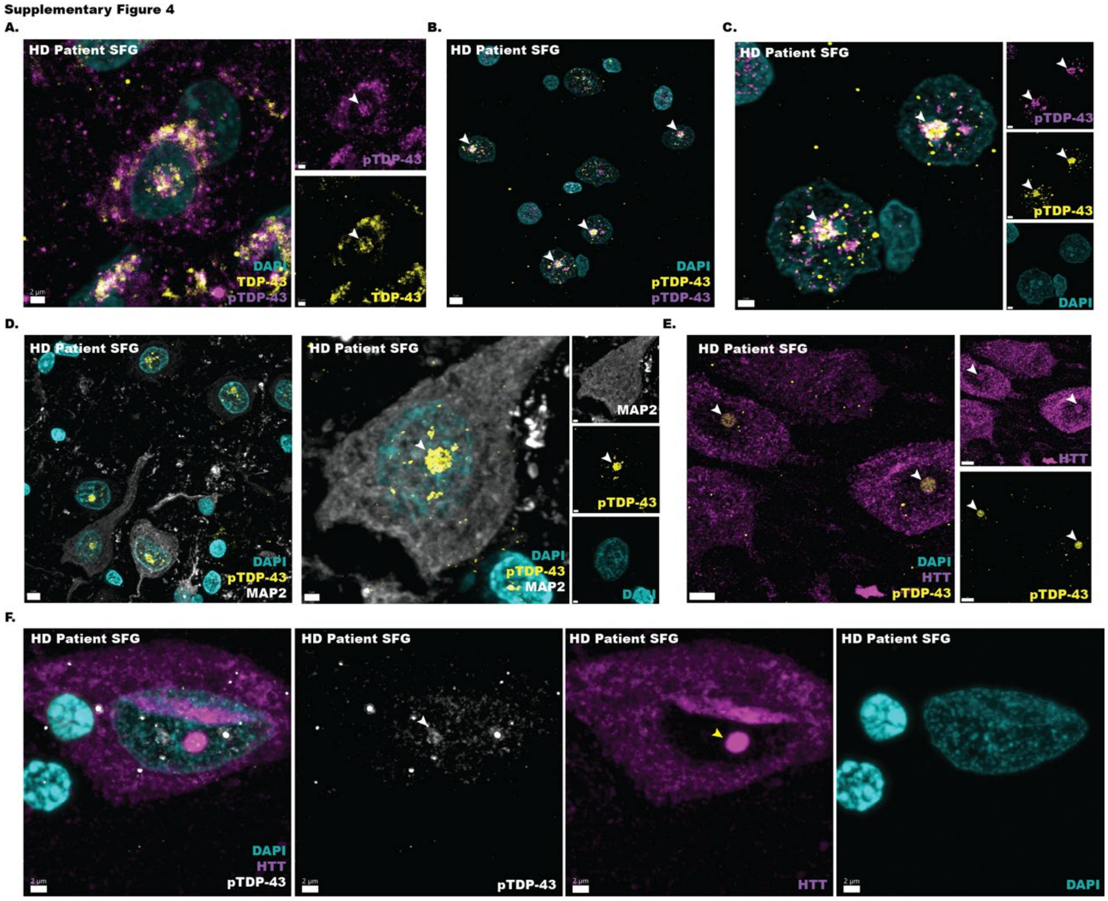


Supplemental Figure 4: (A) Representative images of immunofluorescence (IF) staining showing co-localization of nuclear pTDP-43 AL-bodies (purple) with total TDP-43 (yellow) in the superior frontal gyrus (SFG) of HD patients. White arrows indicate location of AL-bodies. scalebar=2um (B) Lower magnification IF showing the co-localization of two pTDP-43(S409/S410) antibodies: pTDP-43 (purple, Cat#RB3655 (gift from Dr. Leonard Petrucelli) and pTDP-43 (yellow, Cat# Biolegend 50-102-9913). White arrows indicate location of AL-bodies. scalebar=5um (C) Higher magnification IF showing the co-localization of two pTDP-43(S409/S410) antibodies: pTDP-43 (purple) and pTDP-43 (yellow). scalebar=2um (D) Lower magnification scalebar=5um (left) and higher magnification scalebar=2um (right) IF images showing pTDP-43 AL-bodies (yellow) within MAP2 positive neurons (white). (E) IF images showing co-localization of nuclear pTDP-43 AL-bodies (yellow) with the HTT antibody 5526 (purple). scalebar=5um (F) IF images showing co-localization of nuclear pTDP-43 AL-bodies (white) with the HTT antibody 5526 (purple) and the lack of co-localization when canonical nuclear HTT inclusion (yellow arrow) is detected. scalebar=2um.

### Supp Figure 5: TDP-43 eCLIP-seq in HD R6/2 Striatum and Cortex

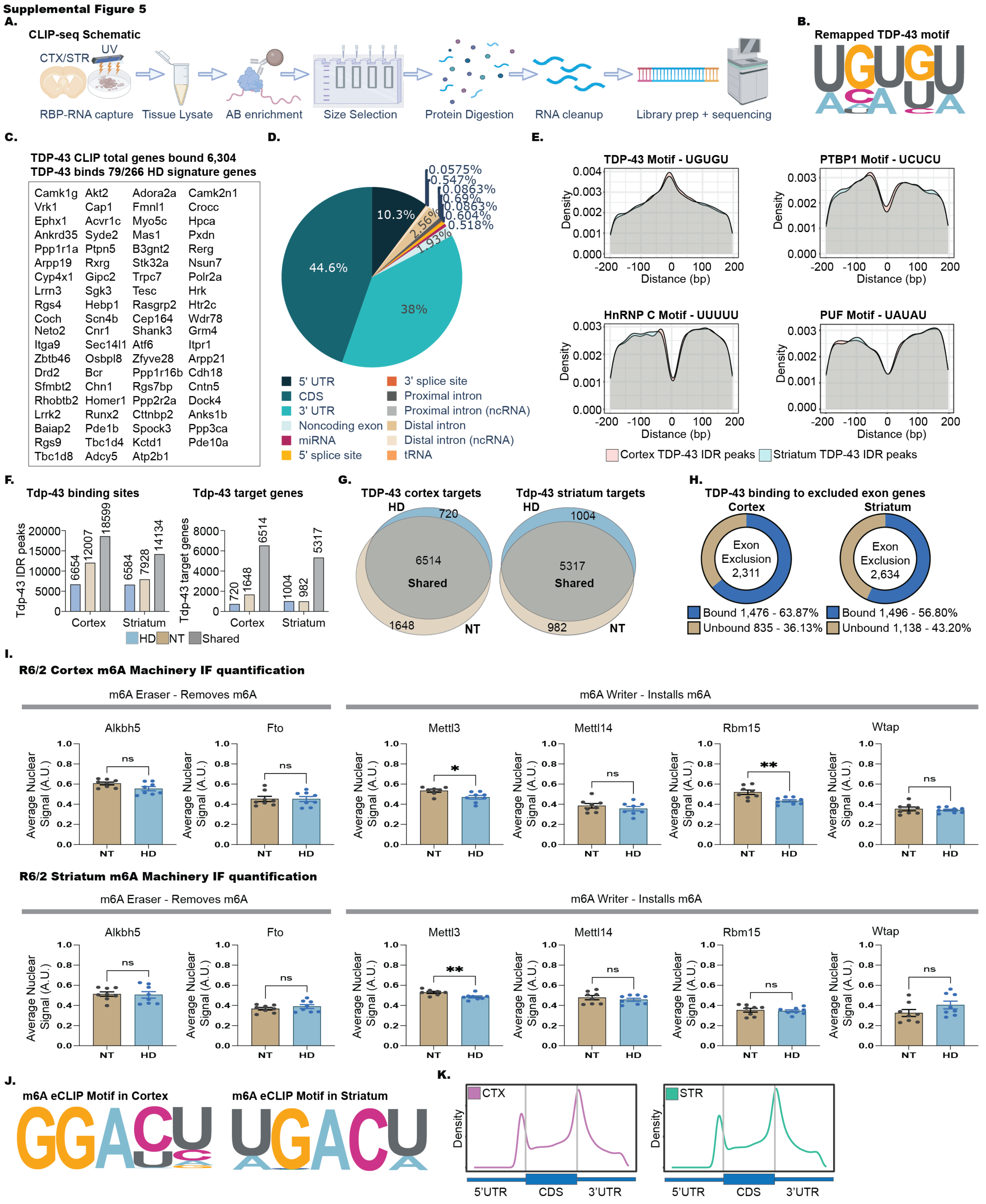

Supp Figure 5: (A) Schematic of TDP-43 CLIP-seq experiment. (B) Enriched motif from Polymenidou et al., 2011 TDP-43 CLIP-seq reanalysis. (C) list of 79/266 HD stritial signature genes that are bound by TDP-43 from TDP-43 CLIP-seq reanalysis. (D) Pie chart showing the classification of TDP-43 peaks in the genome. (E) TDP-43 eCLIP-seq IDR peaks centered at the 0 position. Y-axis represents the density at which the UGUGU, UCUCU, UUUUU, and UAUAU motifs can be found. (F) Bar graph showing the number of TDP-43 CLIP peaks that are significant by IDR and the target genes containing the IDR peaks. For IDR, all male (n=4) TDP-43 CLIP peaks were combined and used to compare to all female (n=4). TDP-43 CLIP IDR peaks. 25,534 HD (7,224 genes), 30,606 NT (8,150 genes) binding sites in the cortex, and 20,942HD (6,313 genes), 22,062 NT (6,281 genes) in the striatum. (G) Venn diagram showing the number of genes that overlaps between HD vs NT TDP-43 CLIP peaks in the cortex and striatum. (H) TDP-43 peaks overlapping with HD exon exclusion (CTX: 1,476/2,311, STR: 1,496/2,634) (1,496/2,634). (I) m6A machinery IF quantification by cellprofiler. Statistical significance determined by unpaired t-test, ns = not significant, * p-value < 0.05, ** p-value <0.01. (J) Enriched motif for Cortex and Striatum m6A-seq IDR peaks. (K) Metagene plot generated by metaplotR showing the distribution of m6A sites across a metagene. Note: the peak at the 5’UTR has not been filtered for the RNA modification m6Am which is also picked up by the m6A antibody.
